# BoltzProt-1: Towards Efficient De Novo Binder Design with Good Developability

**DOI:** 10.64898/2026.06.23.733997

**Authors:** Talip Uçar, Jack Bates, Yunguan Fu, Wenxian Shi, Hannes Stark, Demitri Nava, Luca Cavalleri, Jeremy Wohlwend, Gabriele Corso, Saro Passaro

## Abstract

Designing binders against novel protein targets remains a central challenge in computational drug discovery. Here we introduce **BoltzProt-1**, a pipeline for generating protein binders, including nanobodies, with improved hit rates and favorable developability properties. At its core lie a refined iteration of BoltzGen’s generative model and a novel protein-protein interaction prediction model, **BoltzPPI**. Employing BoltzPPI instead of BoltzGen’s standard structure-prediction confidence metrics to rank nanobody (VHH) designs increases the confirmed-binder hit rate from 3.3% to 8.0% across 10 novel targets. Assessed on 10 additional targets used in prior literature, the BoltzProt-1 pipeline obtains nanobody screening hits for 7 of 10 targets, surpassing the 6 of 10 previously reported by Chai-2. Finally, evaluating the developability of BoltzProt-1-designed nanobodies in terms of stability, aggregation, purity, polyspecificity and hydrophobicity reveals that 58% of its confirmed binders pass every criterion, exceeding both BoltzGen (40%) and clinical-stage VHH controls (21%).

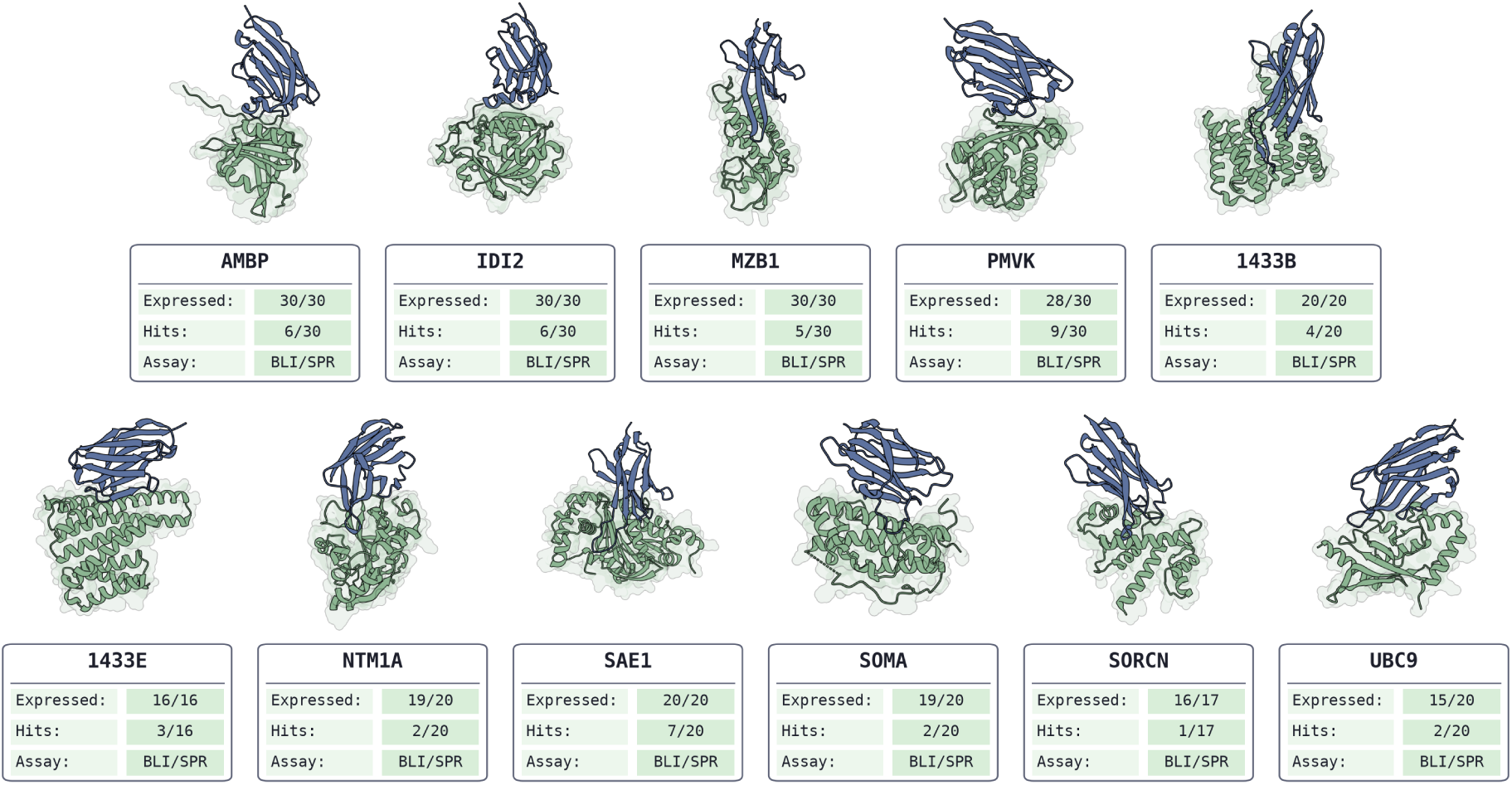

## 1 Introduction

Computational binder design is emerging as an increasingly viable path toward accelerated therapeutic discovery, with several models succeeding on diverse targets [Watson et al., 2023, Bennett et al., 2024, Stark et al., 2025, Zhang et al., 2025]. Realizing the ultimate goal of automating drug discovery requires further expanding the space of accessible targets and improving hit rates. Additionally, designed binders must satisfy the developability criteria required for therapeutic use, such as thermostability and interaction selectivity.

To further approach these desiderata, we introduce the **BoltzProt-1** pipeline, which produces binders (including nanobodies/VHH constructs) against a broader range of targets, with improved per-target hit-rates and robust developability characteristics. At its core is a refined version of the BoltzGen generative model [Stark et al., 2025], combined with a novel protein-protein interaction prediction model **BoltzPPI** for ranking and filtering designs.

The BoltzPPI model translates the insights of Boltz-2 [Passaro et al., 2025] for protein-ligand affinity prediction to protein binder scoring. Ranking novel designs demands generalization to previously unseen chemical matter, which in turn requires that inferences rest on broadly transferable signals such as 3D structural patterns. BoltzPPI therefore grounds its predictions in the pairwise interaction features of a structure prediction model and the 3D coordinates it produces. To further promote generalization, these representations are restricted to a subset concentrated on the predicted structure’s interaction interface. During training, its binary interaction prediction is supervised with protein-protein complexes from the PDB and patent-derived complexes as sources of high-confidence positive labels, alongside synthetically generated protein-protein pairs serving as negatives. The resulting binding-confidence score complements the standard confidence metrics of structure prediction models and proves effective at ranking binder designs.

We evaluate BoltzProt-1 along two axes: binding and developability. For binding, we adopt a more stringent assessment protocol that distinguishes initial screening hits (what prior binder design model literature typically reports as binders) from confirmed binders (see Section 3.1). Under this classification, BoltzPPI raises the confirmed-binder hit rate from 3.3% to 8.0% when designing nanobodies against a panel of 10 novel targets that are low-similarity to any bound complex in the training data. Evaluated on 10 targets from Chai-2 [Chai et al., 2025], BoltzProt-1 yields screening hits against 7 of 10 targets for nanobody design, surpassing the 6 of 10 reported by Chai-2. We further test the developability properties of BoltzProt-1-designed nanobodies in assays spanning thermal stability, aggregation, monomer purity, hydrophobicity, nonspecific binding, and self-association. Across every category, our designs either match or exceed clinical-stage nanobody controls, and 58% pass all developability criteria simultaneously, compared with 40% of BoltzGen-selected binders and 25% and 21% of the clinical-stage IgG and VHH controls, respectively.

We provide the Boltz API and the Boltz Lab platform with a graphical user interface (see Appendix E) as easy-to-use access points, complete with a model-context protocol for interfacing with large language models. By offering these efficiency-optimized endpoints, we hope to place state-of-the-art binder design within affordable reach of any researcher.

## 2 Methods

### 2.1 BoltzProt-1

BoltzProt-1 is a de novo binder design pipeline that combines an *improved* generative model for candidate generation with **BoltzPPI**, an interaction-aware scoring model. The generative model is an all-atom system that jointly performs binder design and structure prediction, producing large candidate pools. BoltzPPI extends Boltz-2 [Passaro et al., 2025] by adding a protein–protein interaction (PPI) prediction head trained jointly with a confidence head. Given a proposed binder–target complex, the PPI head outputs a binding-confidence score used to select candidates for experimental testing.

These components are combined in a two-stage de novo design pipeline. First, candidate sequences and their predicted binder–target complexes are generated with the generative model. Each candidate complex is then scored with BoltzPPI, and the top-ranked designs are advanced to experimental testing.

### 2.2 BoltzPPI

#### Training

The PPI head is trained jointly with the confidence head on top of the base Boltz-2 representations. It takes as input token-level features, pairwise features, predicted coordinates, minimum inter-token distances, and binder/target token masks. Pairwise PPI inputs are constructed from three components: projected single-token features expanded into pair space, binder-specific mask embeddings indicating the two binding partners, and discretized distance embeddings derived from both the full predicted coordinate distance matrix and the minimum inter-token distance matrix. The single-token representation is augmented with corresponding binder-mask embeddings. These single and pair representations are then refined by a dedicated Pairformer stack comprising 4 blocks, each with 16 attention heads and a dropout rate of 0.25.

During training, the module is exposed to two input regimes. In 50% of cases, it uses both coordinate-derived features and trunk pairwise features. In the remaining 50%, the normalized trunk pairwise representation is dropped. This encourages the model to use geometric information while reducing over-reliance on trunk representations. Together, these regimes define a multi-view training setup in which each view exposes a complementary subset of the input signal [Ucar et al., 2021]. We further regularize training by adding Gaussian noise to randomly selected dimensions of the single and pair representations.

The PPI head is co-trained with the confidence head to jointly predict binary interaction probabilities and structure confidence metrics from the same refined representations. Given an aggregated representation, the PPI head produces a logit *y*^, which is mapped to a probability *p* = *σ*(*y*^). The interaction objective is optimized using focal loss:

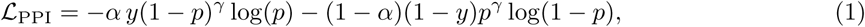

where *y* ∈ {0, 1} is the binding label.

In parallel, the confidence head is trained to predict pLDDT-, PDE-, and PAE-related quantities from the same representations. The two objectives are optimized jointly, with the interaction loss acting as an auxiliary signal:

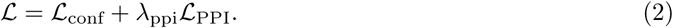

#### Training data

Training data consists of structural protein–protein interaction complexes derived from the PDB [Berman et al., 2000] and patent-derived complexes, alongside synthetically generated protein-protein pairs serving as negatives. This includes antibody–antigen complexes, protein–peptide complexes, and general protein–protein interactions, providing coverage across diverse binding interfaces.

#### Inference

At inference time, BoltzPPI operates on a proposed binder–target complex using the same single-token and pairwise representations, predicted coordinates, and distance-derived features described before. These inputs are refined by the Pairformer stack to produce interaction-aware single and pair representations. The interaction score is computed from two pooled summaries: an average of pair features over inter-chain token pairs and an average of single-token features over all binder and target tokens. After layer normalization, these pooled representations are concatenated, processed by a multilayer perceptron, and mapped to a scalar probability via a sigmoid output to produce the interaction score.

The same refined representations are also used to compute confidence metrics, including pLDDT, PDE, and PAE, together with aggregate complex- and interface-level metrics. In the full design pipeline, the binding confidence is the primary ranking signal for selecting candidates for experimental validation.

## 3 Binding Results

### 3.1 Binding classification

In an assay such as surface plasmon resonance (SPR) or biolayer interferometry (BLI), the designed protein is attached to a surface, and a solution containing the target is flowed across it, while recording a signal that tracks how the surface changes over time. This results in a set of curves called a sensorgram. In an ideal case, a design that does not bind gives a flat line, whereas for a binder, a clear rise and fall is present, and a set of equations can be fit to the curves to compute an affinity. In practice, other effects besides one-to-one design-target interaction can also produce a signal or noise in the sensorgram^1^.

As a consequence, a primary screen often yields sensorgrams that indicate an interaction but do not confidently confirm binding between the binder and the target. Much of the literature reports such cases as binders. We instead adopt the following categorization:

- **Screening hit:** The sensorgrams indicate an interaction. This could be a clean sensorgram with clear binding, but it also includes sensorgrams with an ambiguous signal. The latter case is reported as a hit from an initial screen, not as a confirmed binder.
- **Confirmed binder:** A design is called a binder only when a clean sensorgram, or similarly conclusive data, confirms binding, and an affinity is reported only when the kinetic data can support one. Such evidence may arise from a first screen, but ambiguous cases require follow-up assays. For instance, varying the capture level of the binder on the plate, adjusting the target’s concentration range, or inverting the assay orientation as to whether the target or the design is attached to the plate.

The binding experiments are carried out by Adaptyv Bio and Sino Biological. In our experiments, the **confirmed binders** are validated in an orthogonal binding assay, providing increased confidence that binding is not specific to the initial assay format. We use a *flipped* assay in which the immobilization orientation is reversed to reduce format-specific and avidity artifacts as confirmation. In the BoltzGen low-homology target panel, screening hits from both BoltzGen and BoltzProt-1 are also evaluated by Sino Biological for independent binding testing. For the low-homology target panel, we report screening hits and confirmed binders separately. For the ten-target Chai-2 panel (Section 3.3), we report screening hits. We provide the sequences of all the binders in Appendix D.1 (Table 4) together with their BLI/SPR sensorgrams, including the flipped-format confirmation assay, in Appendix D.2.

**Table 1:** Binding summary on the low-homology target panel. BoltzGen and BoltzProt-1 each test 15 designs per target from the same BoltzGen candidate pools; BoltzProt-1 selections exclude sequences already selected by BoltzGen. “Hits” = initial screening hits; “Revised Hits” = screening hits after the revised Adaptyv protocol; “Binders” = confirmed binders. Screening-hit rates are computed from the revised screening-hit counts. Target success is the number of targets with at least one confirmed binder.

| Method | Designs | Hits | Revised Hits | Hit rate | Binders | Binder rate |
| --- | --- | --- | --- | --- | --- | --- |
| BoltzGen | 150 | 15 | 7* | 4.7% | 5 | 3.3% |
| BoltzProt-1 | 150 | 24 | 14* | 9.3% | 12 | **8.0%** |

\*Screening-hit counts decreased after Adaptyv revised its screening-hit calling protocol.

### 3.2 BoltzProt-1 improves nanobody hit rate on novel targets

We first evaluate BoltzProt-1 on the BoltzGen low-homology target panel, which is designed to test generalization to targets without close bound homologs in the PDB [Stark et al., 2025]. The panel comprises AMBP, GM2A, HNMT, IDI2, METTL16, MZB1, ORM2, PHYH, PMVK, and RFK. To isolate the effect of ranking, we compare BoltzProt-1 to the published BoltzGen selections using the same candidate pools and testing budget. Both methods test 15 candidates per target, 150 nanobody designs across the 10 low-homology targets, drawn from the same published BoltzGen candidate pools; BoltzPPI’s selections exclude sequences already chosen by BoltzGen. By holding the candidate pool fixed and varying only the selection, this setup directly tests whether BoltzPPI improves recovery from the available designs. BoltzPPI recovers **24 screening hits** (**16.0%**) and **12 confirmed binders** (**8.0%**), compared with **14 screening hits** (**9.3%**) and **5 confirmed binders** (**3.3%**) for BoltzGen, a 2.4-fold improvement in confirmed-binder rate.

BoltzPPI recovers confirmed binders for **3 of 10** targets (AMBP, IDI2, PMVK), compared with **2 of 10** for BoltzGen (IDI2, PMVK), improving recovery on targets. With an optimized generative model, BoltzProt-1 is additionally able to recover a confirmed binder for MZB1, extending the target success rate to **4 of 10** targets (AMBP, IDI2, PMVK, MZB1). Further confirmation of the full set of *screening hits* is ongoing.

As an additional external reference, we compare BoltzProt-1 with Protenix-v2 [Zhang et al., 2025] on the two targets for which they report experimental results (Figure 1b). The protocols are not identical: Protenix-v2 reports larger experimental test sets per target, with 2 hits out of 50 designs for one AMBP epitope, 13 out of 27 for another AMBP epitope, and 12 out of 50 for IDI2. At the screening-hit level, BoltzProt-1 achieves hit rates comparable to or higher than Protenix-v2 on both AMBP and IDI2.

**Figure 1:**
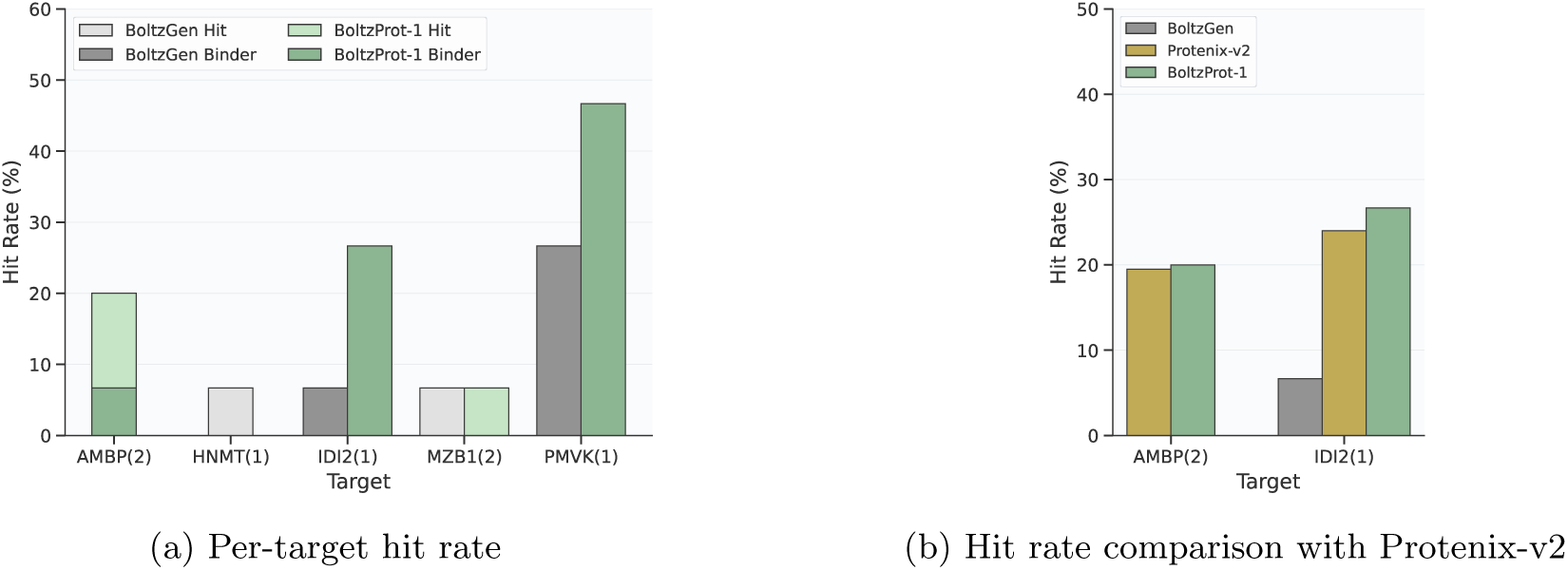
Design hit rate comparison on BoltzGen low-homology targets. **a)** Per-target screening-hit (lighter bar) and confirmed-binder (darker bar) rates for BoltzGen and BoltzProt-1 on the low-homology target panel. **b)** Comparison of screening hit rates with Protenix-v2 on the two common targets (AMBP, IDI2). Assay valency is marked in brackets.

### 3.3 Nanobody design on Chai-2 panel targets

To test whether BoltzProt-1 transfers beyond the BoltzGen low-homology target panel, we evaluate it on the ten protein targets reported in the Chai-2 antibody-design study [Chai et al., 2025]. The set includes signalling and adaptor proteins (14-3-3*β*/*α* [1433B], 14-3-3*ε* [1433E]), secreted hormones and cytokines (leptin [LEP], oncostatin-M [ONCM], somatotropin [SOMA]), calcium-binding and regulatory proteins (S100-B [S100B], sorcin [SORCN]), and enzymes involved in SUMOylation and protein methylation (SUMO-activating enzyme subunit 1 [SAE1], SUMO-conjugating enzyme UBC9 [UBC9], and the N-terminal methyltransferase NTMT1 [NTM1A]). More information about the targets is provided in Appendix D.1.

On this panel, BoltzGen and BoltzProt-1 are run as separate end-to-end design pipelines, with each method generating and selecting its own candidates, and we include the published Chai-2 VHH results as an external comparison [Chai et al., 2025]. All methods test up to 20 sequences per target (Chai-2 does not always reach this budget). BoltzGen generates 60k candidates per target and BoltzProt-1 generates 240k, from which the top-ranked designs are selected for testing. Because the three methods differ in design setup, generation budget, and candidate selection, this experiment compares full pipelines rather than isolating the effect of ranking within a fixed candidate pool. We report *screening hits*, with further confirmation ongoing.

#### BoltzProt-1 achieves the broadest target coverage

BoltzProt-1 recovers screening hits for **7 of 10** targets (1433B, 1433E, NTM1A, SAE1, SOMA, SORCN, and UBC9), more than any other method, compared with **6 of 10** for Chai-2 and **3 of 10** for BoltzGen (Figure 2b). Relative to BoltzGen, which finds hits only on 1433B, NTM1A, and S100B, BoltzProt-1 retains 1433B and NTM1A and adds five further targets (1433E, SAE1, SOMA, SORCN, and UBC9); it does not recover S100B, the one target where BoltzGen succeeds but BoltzProt-1 does not. This broad coverage is consistent with the results of Section 3.2: combining an improved generative model with BoltzPPI prioritization recovers experimentally testable binders across a more diverse set of targets.

**Figure 2:**
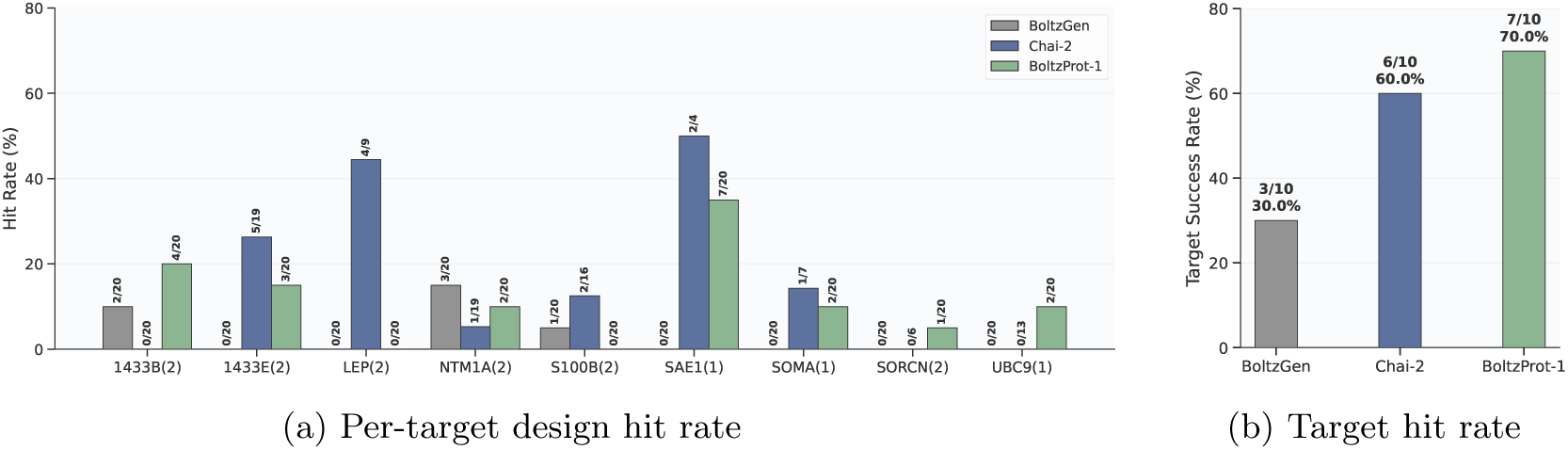
Nanobody design on the Chai-2 VHH targets. **a)** Screening hit ratio per target. **b)** Fraction of targets with at least one screening hit. Chai-2 numbers are the published VHH results [Chai et al., 2025]. Assay valency is marked in brackets.

### 3.4 Prior structural coverage of target binding sites

We also assessed whether the target binding sites used by our designs resemble protein–protein interfaces already observed in the PDB. Using FoldDiSCO [Kim et al., 2026], we compare these binding-site geometries to known bound-interface geometries and summarize the result by the best binding-site similarity score and the number of matching bound-interface homologs (Figure 3). The low-homology target panel shows low similarity to known bound interfaces and few matching homologs, consistent with the original motivation of the BoltzGen benchmark. The Chai-2 panel spans a broader range, with several targets showing higher similarity to known PDB interfaces. This analysis provides context for interpreting the experiments: the low-homology target panel emphasizes binding sites with limited prior structural precedent, whereas the Chai-2 panel includes targets with more variable bound-interface coverage.

**Figure 3:**
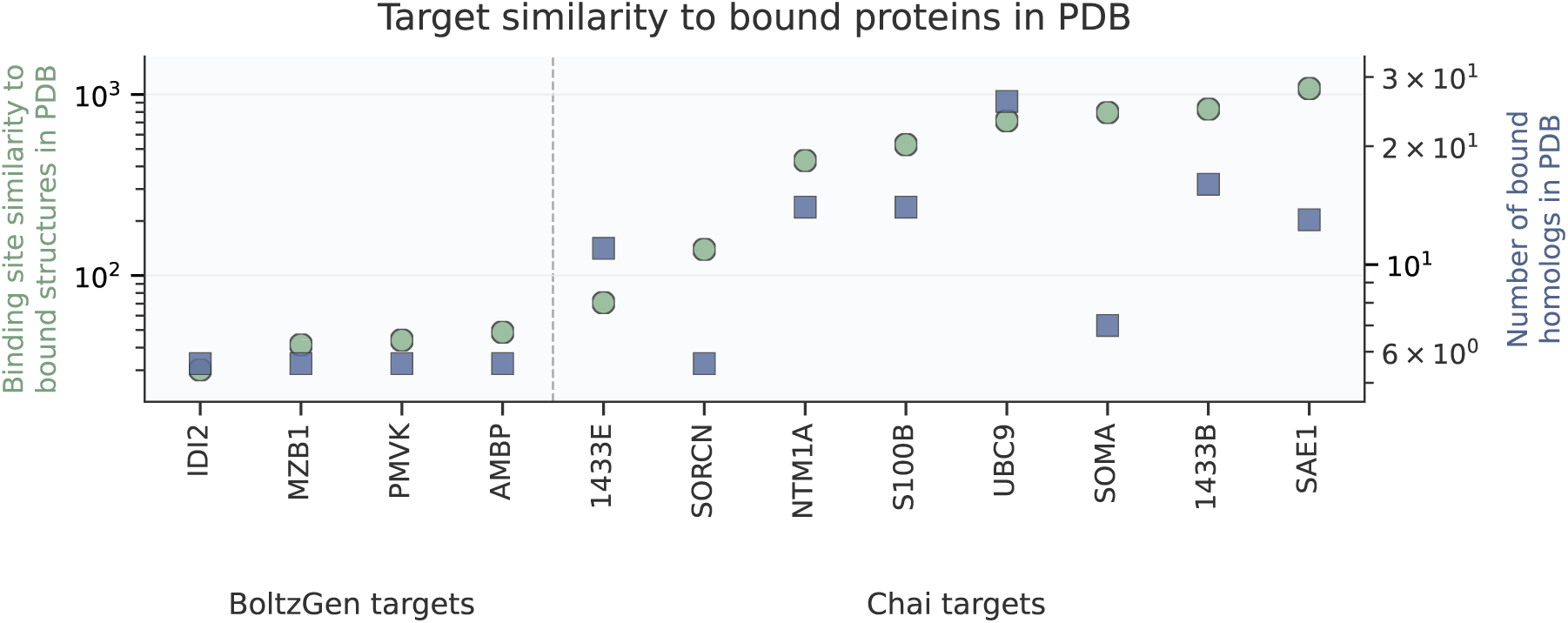
Target similarity to bound proteins in the PDB. For each target, we report binding-site similarity to proteins observed in bound PDB structures and the number of bound homologs in the PDB. The low-homology target panel has low similarity and few bound homologs, whereas the Chai-2 panel spans a wider range and includes several targets with substantially more bound structural precedent.

### 3.5 Recovered binders are distant from known SAbDab CDRs

We next ask whether the experimentally recovered designs closely resemble known antibody or nanobody sequences. For each BoltzProt-1 confirmed binder from the low-homology target experiments and each BoltzProt-1 screening hit from the Chai-2 target panel, we compute the minimum edit distance to sequences in SAbDab[Dunbar et al., 2014], using a database snapshot with cutoff date May 27, 2026. We compute distances both over the concatenated CDR1, CDR2, and CDR3 sequences and over CDR3 alone.

Across both target panels, every recovered design has a minimum CDR3 edit distance of at least four to its closest SAbDab match, with larger minimum edit distances when CDR1, CDR2, and CDR3 are considered together (Figure 4). This pattern holds for confirmed low-homology target binders as well as screening hits on the Chai-2 panel, indicating that the recovered binders are not simple matches to known antibody or nanobody CDRs. This analysis provides a sequence-level check complementary to the structural target-similarity analysis above: BoltzProt-1 recovers experimentally active binders whose CDRs remain distinct from known SAbDab entries.

**Figure 4:**
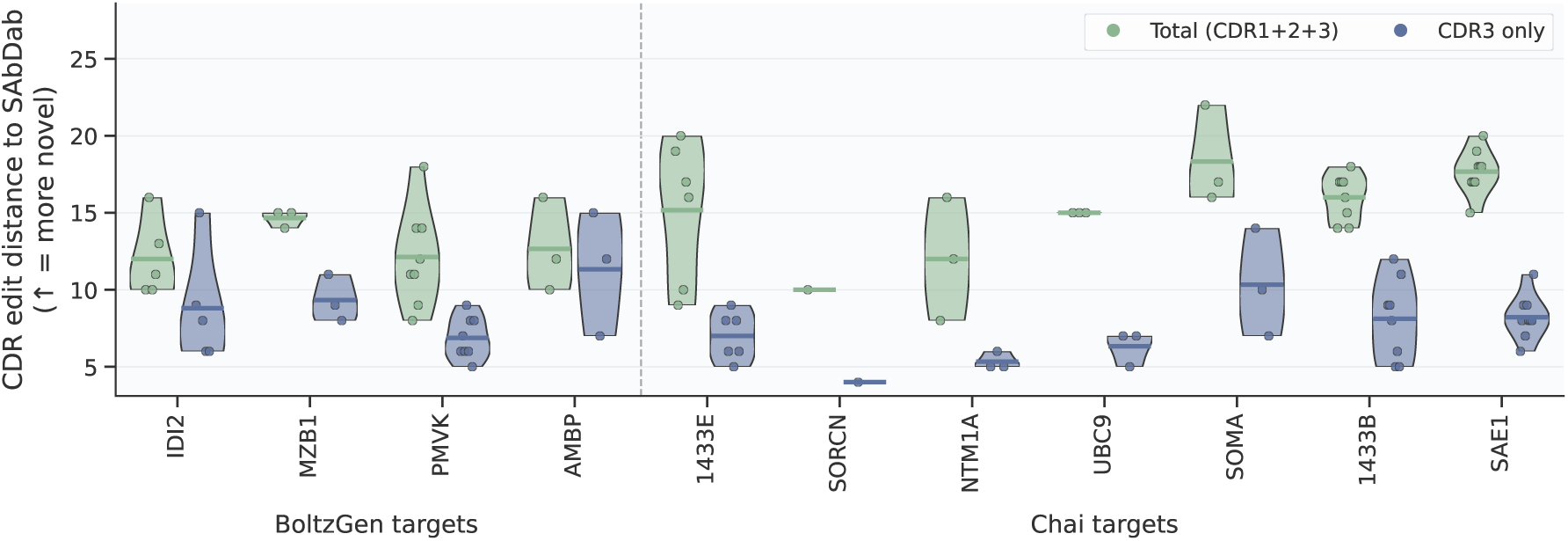
CDR edit distance to known SAbDab antibodies and nanobodies. For each BoltzProt-1 confirmed binder from the low-homology target experiments and each BoltzProt-1 screening hit from the Chai-2 target panel, we compute the minimum edit distance to antibody and nanobody sequences in SAbDab using a database cutoff date of May 27, 2026. Distances are shown for concatenated CDR1+CDR2+CDR3 sequences and for CDR3 alone. Higher values indicate greater sequence distance from the nearest known SAbDab entry.

## 4 Developability Results

### 4.1 Developability assays

A binder is only useful as a therapeutic if it can be expressed, purified, formulated, and manufactured at scale, which requires that it remain stable, soluble, and monomeric while avoiding nonspecific and self-interaction liabilities. These developability properties often determine whether a high-affinity binder can advance at all, so we evaluate the BoltzGen and BoltzProt-1 designs from the first low-homology target experiment across a comprehensive developability panel, with all experiments carried out by Twist Bioscience (see Appendix Section C for details). Each assay probes a distinct liability: thermal stability (*T*_onset_, *T*_m1_, *T*_m2_) and aggregation onset (*T*_agg_) report folding stability and resistance to aggregation; monomer purity by analytical size-exclusion chromatography (aSEC) and sample heterogeneity via the polydispersity index (PDI) report homogeneity; surface hydrophobicity by hydrophobic interaction chromatography (HIC) reports a key driver of aggregation and nonspecific binding; nonspecific binding by baculovirus particle ELISA (BVP) reports polyspecificity; and self-association propensity by affinity-capture self-interaction nanoparticle spectroscopy (AC-SINS) reports the tendency to self-associate at high concentration. Following prior work [Chai et al., 2025], developability is assessed relative to control molecules and across the full assay panel rather than through independent per-assay thresholds alone. The control set comprises clinical-stage IgG antibodies and VHH-Fc constructs, with VHH controls aligned to those used in [Chai et al., 2025]; all designed VHHs are evaluated in VHH-Fc format, whereas antibody controls are measured in IgG format. The full list of controls is provided in Table 2 in Appendix C.1.

### 4.2 BoltzProt-1 binders exhibit good developability

We evaluate developability on the confirmed binders from the low-homology target experiment (12 for BoltzProt-1, 5 for BoltzGen) relative to 36 clinical-stage controls (12 IgG and 24 VHH). Across the thermal panel, BoltzProt-1-selected binders match BoltzGen-selected binders and VHH controls on *T*_m1_, *T*_m2_, and *T*_agg_ (Figure 7a–d), indicating stable folding and delayed aggregation onset, and they perform well across the remaining panel of monomericity, dispersion, hydrophobicity, nonspecific binding, and self-interaction assays (Figure 5). Under the combined criteria, BoltzProt-1 binders achieve the highest overall pass rate of any cohort: 58% pass all filters (7/12), exceeding BoltzGen (40%, 2/5) as well as the clinical-stage controls (IgG: 25%, 3/12; VHH: 21%, 5/24). The cumulative filtering analysis (Figure 6) confirms this robustness, with BoltzProt-1 designs showing minimal early-stage attrition, a single reduction at PDI (12 to 11) followed by full retention through thermal stability and monomer purity, and the only meaningful loss occurring at the hydrophobicity stage (HIC, 11 to 7); failures in the control cohorts are likewise driven mainly by polyspecificity assays rather than by thermal stability or monomer purity. This strength is especially notable against the Chai-2 developability benchmark, which reports a 37% failure rate at the initial monomer-purity gate (≥ 90% purity to advance designs) [Chai et al., 2025]: the BoltzProt-1 confirmed binders clear this gate without loss of monomer purity or related quality, showing that BoltzProt-1’s improved binding recovery is paired with excellent developability. Figure 6 summarizes this high-quality outcome, the fraction of confirmed binders that additionally pass all strict developability filters.

**Figure 5:**
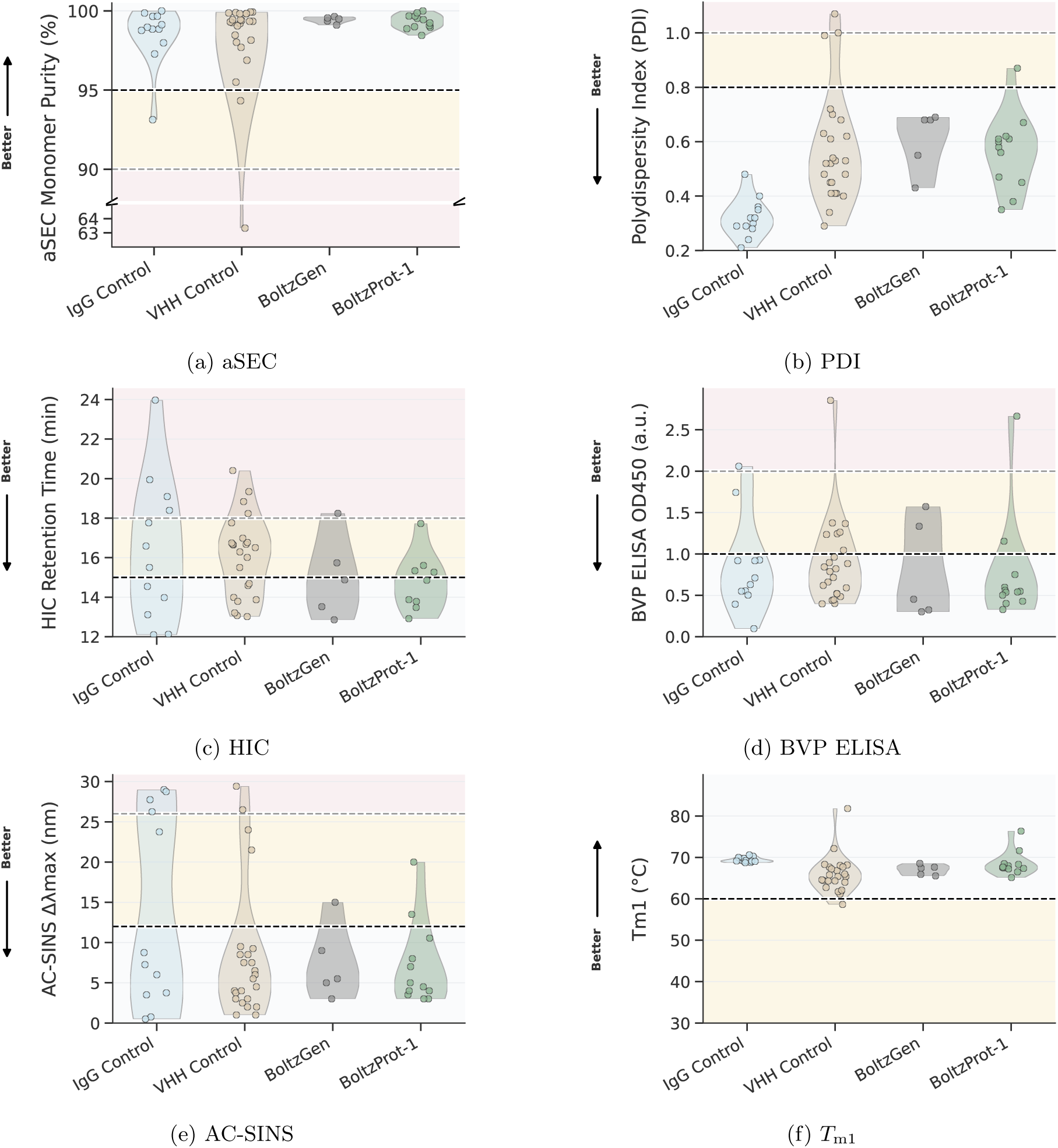
Developability assays. BoltzProt-1 binders across monomericity, dispersion, hydrophobicity, nonspecific binding, self-interaction, and thermal stability assays.

**Figure 6:**
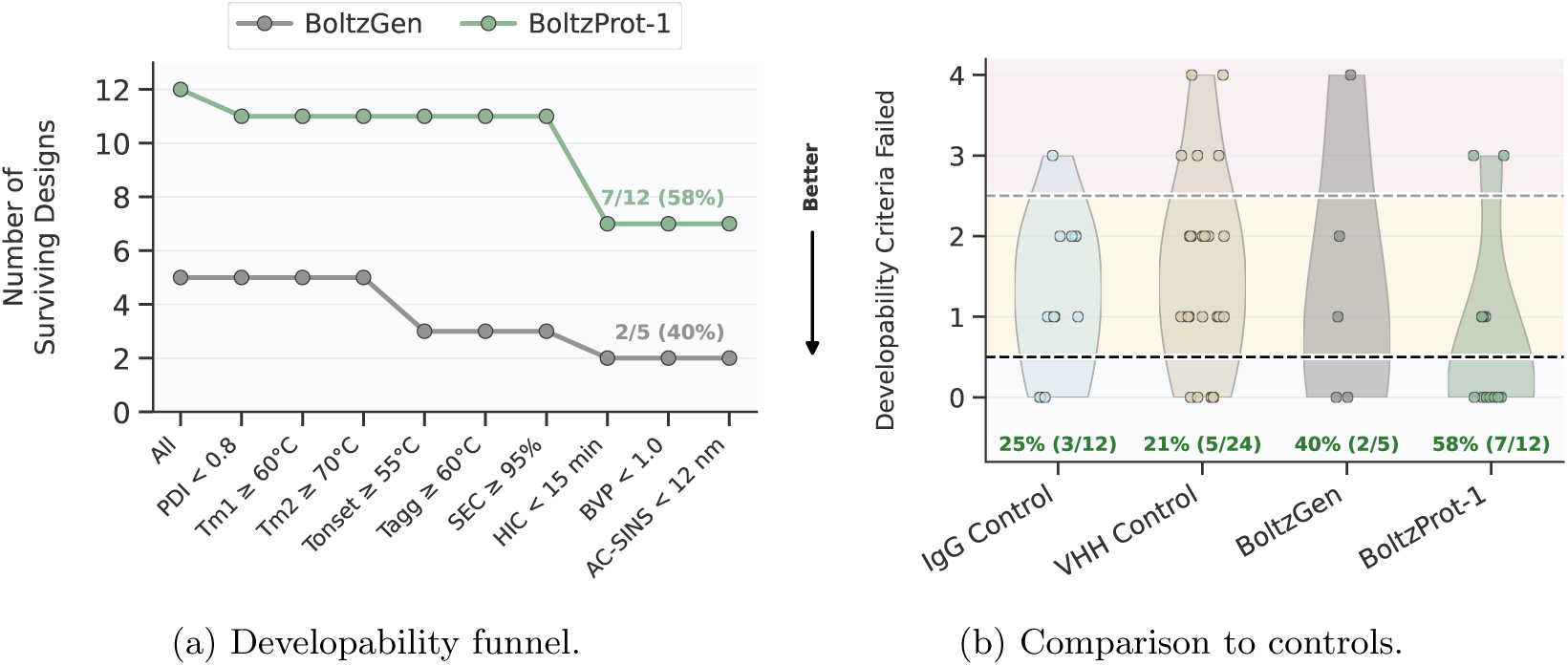
Developability funnel. 58% (7 of 12) of the confirmed BoltzProt-1 binders satisfy all developability criteria, compared with 40% (2 of 5) of the confirmed BoltzGen binders.

## 5 Conclusion

We introduced **BoltzProt-1**, a de-novo binder design pipeline that pairs an improved BoltzGen-style generative model with **BoltzPPI**, an interaction-aware scoring model built on Boltz-2 that predicts protein–protein binding directly from a proposed complex. By ranking candidates with a signal aligned to experimental binding rather than geometric plausibility alone, BoltzProt-1 selects a small, high-quality subset of designs for testing, addressing the prioritization bottleneck that limits generative binder design.

The approach delivers consistent gains across two prospective nanobody settings. On the BoltzGen low-homology target panel, a budget-matched comparison on the same candidate pools raises the confirmed-binder rate from 3.3% to 8.0%, a 2.4-fold improvement, and doubles confirmed-target coverage from 2 to 4 of 10, demonstrating that better selection alone, without changing the generative model or the experimental budget, substantially improves recovery. On a separate ten-target panel from Chai-2, BoltzProt-1 attains the broadest coverage of any method, recovering screening hits for 7 of 10 targets, ahead of both Chai-2 (6 of 10) and BoltzGen (3 of 10), showing that the gains generalize beyond a single benchmark. Beyond binding, BoltzProt-1 excels at developability: across a comprehensive panel spanning stability, aggregation, purity, polyspecificity and hydrophobicity, its binders achieve the highest pass rate of any cohort, surpassing both BoltzGen-selected designs and clinical-stage controls. The recovered binders are also structurally and sequence-wise novel, with CDRs distant from known SAbDab entries and target interfaces with limited prior precedent in the PDB.

The full BoltzProt-1 pipeline is available through the Boltz API and the Boltz Lab platform, making efficient, developability-aware binder design readily accessible for prospective campaigns.

## Supplementary Material

### A Developability

**Figure 7:**
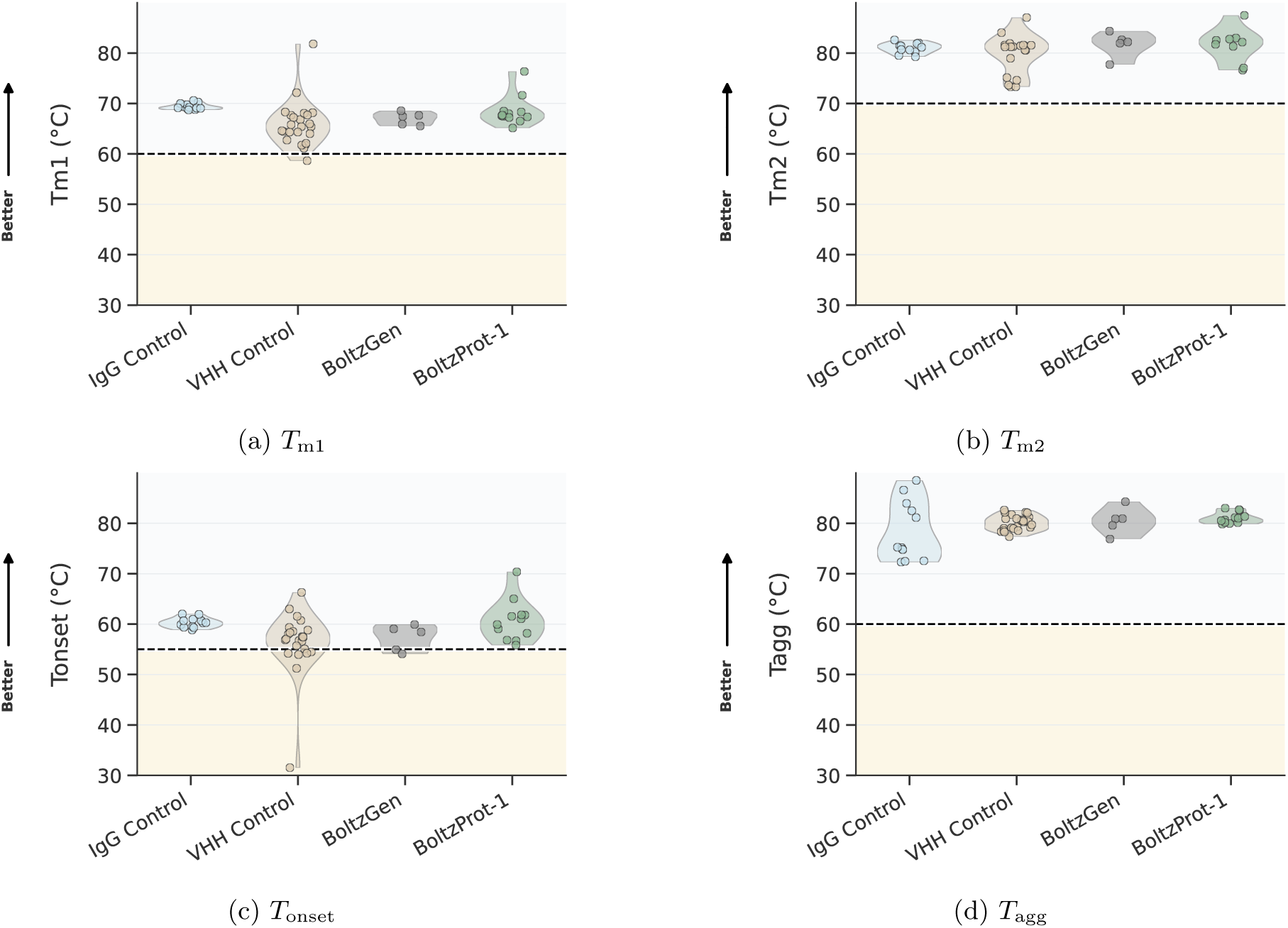
Thermal stability. BoltzProt-1 and BoltzGen binders across *T*_m1_, *T*_m2_, *T*_onset_, and *T*_agg_.

### B Binding Assays

DNA constructs encoding the ligands, each fused to a C-terminal assay tag, were designed by reverse-translating the target protein sequences. The sequences were optimized for manufacturability and yield using codon optimization algorithms to maximize expression efficiency. The optimized DNA constructs of the variants and the 3’ prime fragment containing the linker and affinity tags were ordered as Gene Fragments from Twist Bioscience. Constructs were assembled using the NEBuilder HiFi DNA Assembly Kit (NEB) in 2 *µ*L reactions. The assembled products were characterized by capillary electrophoresis (Agilent ZAG DNA Analyzer, ZAG-135-5000 Kit, FA/ZAG 96-Capillary Array Short, 33cm), and their concentrations were measured using the Qubit DNA Quantification Kit (Invitrogen). Ligand expression was carried out in 8 *µ*L reactions using an optimized prokaryotic in-vitro translation system and 4 nM of assembled gene fragment. Reactions were incubated at 37°C for 8 hours to ensure efficient protein synthesis. Post-expression, protein concentration and yield were normalized using an affinity-based quantification assay.

#### B.1 Biolayer Interferometry (BLI) Affinity Characterization (Adaptyv)

Kinetic binding measurements were performed on a BLI instrument (Gator Bio) using Strep-Tactin XT probes for capture of Twin-Strep–tagged ligands.

##### Sensor Preparation and Capture

Probes were equilibrated in running buffer prior to use. Ligands were captured with the following sequence:

- Baseline 1: 120 s in running buffer
- Ligand loading: 120 s (target loading shift 0.5–1.0 nm)
- Baseline 2: 200 s in running buffer

Running buffer consisted of 10 mM HEPES, 150 mM NaCl, 3 mM EDTA, 0.2% Tween-20, pH 7.4. Temperature was maintained at 25 °C. Data were sampled at 5 Hz.

##### Association/Dissociation (Multi-Cycle Kinetics)

Antigen solutions were prepared as a half-log dilution series (1,000 nM to 30 nM, 4 concentrations) in running buffer.

Each cycle consisted of:

- Association: 220 s in antigen solution
- Dissociation: 240 s in running buffer

Between cycles, probes were regenerated in 10 mM Glycine-HCl, pH 1.5, applied five times for 10 s each, followed by neutralization in running buffer. Negative controls (buffer-only and non-binding ligand) were included for reference subtraction and drift correction.

##### Data Processing and Model Fitting

Data was processed and fitted globally to a 1:1 Langmuir binding model using Adaptyv Fitting software. In summary, all sensor grams underwent standardized preprocessing and curve fitting to ensure accurate kinetic parameter estimation. Preprocessing included trimming to relevant phases (association, dissociation, baseline), correcting signal jumps at transitions, aligning association and dissociation phases, and subtracting baseline and reference signals.

Fitting proceeded through multiple methods in a prioritized order. Initial fits were performed individually using global fitting followed by full, dissociation-only, or slope-based models, if this was not possible group-level models such as equilibrium (saturation), constant (flat), and semi-log linear (linear) were applied. Final kinetic parameters (kon, koff, *K_D_*) were selected based on fit quality. Global fitting was applied using a 1:1 model across all concentrations. In global mode, koff and *K_D_* were fitted directly, and kon was calculated as koff / *K_D_*. Fits were scored and filtered based on quality metrics.

Ligands were classified as binders (True) or non-binders (False) based on the presence of quantifiable binding curves and calculated *K_D_* values. In cases where ligands produced a significant signal shift during the association phase (≥ 300% over the negative control), but the signal could not be reliably fit, binding labels were assigned based on the magnitude of the observed shift.

#### B.2 Surface Plasmon Resonance (SPR) Affinity Characterization (Adaptyv)

Binding kinetics of the expressed ligands to their cognate targets were characterized using Surface Plasmon Resonance (SPR) on a Carterra LSA XT instrument. Briefly, twin-strep-tagged ligands were captured on a sensor surface functionalized with Strep-Tactin XT (IBA Lifesciences), followed by sequential injections of increasing antigen concentrations.

##### Chip Preparation (Strep-Tactin XT Immobilization)

Surface functionalization of the carboxymethylated chip was performed following a multi-step chip preparation protocol: Conditioning - The sensor surface was conditioned with NaOH (50 mM) to remove residual contaminants and prepare the surface for activation. Activation - The surface was activated using freshly prepared EDC/NHS (1-Ethyl-3-(3-dimethylaminopropyl)carbodiimide/N-Hydroxysulfosuccinimide) (Xantec) solution to form reactive esters. Capture - Strep-Tactin XT was diluted to 50 *µ*g/mL in 10 mM sodium acetate buffer (pH 4.5) and injected onto the activated surface to allow covalent coupling. Quenching - Excess reactive groups were quenched with 1 M ethanolamine hydrochloride (pH 8.5). Wash - The surface was washed with 0.1 M sodium borate, 1 M NaCl pH 9.0 to remove loosely bound material. Reference and capture sensors underwent identical treatment, allowing for accurate background subtraction.

##### Ligand Capture and Kinetics

Following surface preparation, ligands containing a twin-strep-tag were captured via specific interactions with immobilized Strep-Tactin XT using the multichannel head (96 channel array) using bidirectional flow for 750s followed by a baseline step in running buffer for 600s. The analyte was diluted in running buffer (10 mM HEPES, 150 mM NaCl, 3 mM EDTA, 0.05% Tween-20, pH 7.4) to 7 concentrations in a half-log dilution series (e.g., 1000 nM to 1 nM). Each cycle is repeated over increasing concentration for the analyte to generate the single cycle kinetic assays without intermediate regeneration at a flow rate of 50*µ*L/min. Each assay cycle consisted of: Baseline phase: 60 seconds of buffer-only flow Association phase: 300 seconds of analyte injection Dissociation phase: 600 seconds of buffer-only flow Following each full series of interactions, the chip surface was regenerated using 10 mM glycine-HCl (pH 1.5) for 5 mins, followed by wash in running buffer for 20 mins.

##### Data Processing and Analysis

Data was processed and fitted globally to a 1:1 Langmuir binding model using Adaptyv Fitting software. In summary, all sensor grams underwent standardized pre-processing and curve fitting to ensure accurate kinetic parameter estimation. Preprocessing included trimming to relevant phases (association, dissociation, baseline), correcting signal jumps at transitions, aligning association and dissociation phases, and subtracting baseline and reference signals. Fitting proceeded through multiple methods in a prioritized order. Initial fits were performed individually using global fitting followed by full, dissociation-only, or slope-based models, if this was not possible group-level models such as equilibrium (saturation), constant (flat), and semi-log linear (linear) were applied. Final kinetic parameters (kon, koff, *K_D_*) were selected based on fit quality. Global fitting was applied using a 1:1 model across all concentrations. In global mode, koff and *K_D_* were fitted directly, and kon was calculated as koff / *K_D_*. Fits were scored and filtered based on quality metrics. Ligands were classified as binders (True) or non-binders (False) based on the presence of quantifiable binding curves and calculated *K_D_* values. In cases where ligands produced a significant signal shift during the association phase (≥300% over the negative control), but the signal could not be reliably fit, binding labels were assigned based on the magnitude of the observed shift.

#### B.3 Surface Plasmon Resonance (SPR) Affinity Characterization (Sino)

SPR characterisation was performed using a Cytiva Biacore 8K instrument. In each instance, binders were used as the ligand in multi-cycle kinetics runs with the target as analyte. Analytes were diluted into running buffer, and five concentrations were tested, with a top concentration of 1 µM and two-fold dilutions thereof, in addition to a blank. Both association and dissociation were performed for 60 seconds at a flow rate of 30 µL/min.

##### Fc-fusion binders (Protein A capture)

Binders with Fc fusions were captured on a Series S Protein A chip (29127556, Cytiva). Binders were diluted to 5 µg/mL and injected over the flow cell for 1 minute at 10 µL/min. HBS-ET was used as the running buffer. Regeneration was performed using 10 mM glycine-HCl (pH 1.5), injected for 60 s at 30 µL/min.

##### His-tagged binders (NTA capture)

His-tagged binders were captured on an NTA chip (28995043, Cytiva). The surface was first prepared with a one-minute pulse of nickel solution over the active flow cell at a flow rate of 10 µL/min. Binders were then diluted to 1.5 µg/mL in running buffer and injected for 10 seconds at a flow rate of 10 µL/min for capture. 2× HBS-T was used as the running buffer. Regeneration used 350 mM EDTA, injected for 60 s at 30 µL/min.

##### Data Processing and Analysis

Kinetic parameters were determined by fitting a 1:1 Langmuir model to the data. A steady-state model was additionally applied where fast association/dissociation was observed.

### C Developability Assays

Developability experiments were performed by Twist Bioscience using standardized protocols for antibody production, purification, and biophysical characterization. The assays provide a comprehensive assessment of developability, capturing thermal stability, aggregation propensity, self-interaction, hydrophobicity, nonspecific binding, and sample heterogeneity.

#### C.1 List of controls used in developability assays

**Table 2:** Control molecules used for developability assessment. VHH controls are evaluated in VHH-Fc format, and antibody controls in IgG format.

| IgG controls |  | VHH-Fc controls |
| --- | --- | --- |
| Adalimumab | Brivekimig1_VHH1 | brivekimig2_VHH1 |
| Atezolizumab | Erfonrilimab_VHH2 | caplacizumab_VHH1 |
| Dacetuzumab | Gocatamig2_VHH1 | envafolimab_VHH1 |
| Golimumab | Gontivimab_VHH1 | gacovetug_VHH1 |
| Ixekizumab | Isecarosmab_VHH1 | gefurulimab_VHH1 |
| Simtuzumab | Lofacimig_VHH2 | letolizumab_VHH1 |
| Cetuximab | Lunsekimig2_VHH1 | lofacimig_VHH1 |
| Cixutumumab | Lunsekimig3_VHH1 | ozoralizumab_VHH1 |
| Motavizumab | Podentamig1_VHH2 | ozoralizumab_VHH2 |
| Palivizumab | Produvofusp_VHH1 | rimteravimab_VHH1 |
| Pembrolizumab | Sonelokimab1_VHH1 | sonelokimab2_VHH1 |
| Robatumumab | Vobarilizumab_VHH1 | tarperprumig_VHH2 |

#### C.2 Antibody Production and Purification

Twist utilizes the Thermo Fisher Expi293™ Expression System Kit to carry out a lipid-based transfection with its DNA on an Expi293 cell line. After 1 day of incubation at 37 °C, Expi293 Enhancers 1 and 2 are added to the cell plate, which is then incubated at 37 °C for an additional 3 days.

Following incubation, the cells are pelleted by centrifugation and the supernatant is extracted and filtered through a 0.2 µm filter plate. The supernatant is transferred into a 2 mL deep well block with Protein A binding antibody purification resin. The supernatant-resin plate is then mixed in an incubator shaker to allow time for the expressed material to bind.

The resin-supernatant mixture is transferred into a filter plate (20 µm glass fiber membrane). The supernatant is filtered through in a centrifuge and the filter is washed twice by adding DPBS pH 7.4 and centrifuging the filter plate stacked onto a collection plate.

Antibodies are eluted off the resin with either 50 mM citrate (pH 3.0) or 0.1 M glycine (pH 2.7) by centrifuging the filter stacked on a destination plate pre-filled with the full volume of neutralization buffer (1 M HEPES pH 8.0 for citrate, or 1 M Tris-HCl pH 7.5 for glycine, respectively). The elution step is repeated to dispense the second half of the elution buffer, and the purification plate is sealed, mixed on a shaker, and proceeds to quantification.

#### C.3 Analytical Size Exclusion Chromatography (aSEC)

Size-exclusion chromatography (SEC) was used to evaluate the molecular size distribution and structural integrity of protein samples, including detection of monomer content, aggregation, and degradation.

1.5 µg or less (10 µL max injection, injection volume dictated by sample concentration) of each sample was injected onto a MAbPac™ SEC-1 column (4 × 150 mm, 5 µm particle size, 300 Å pore size; Lifetech) on a Vanquish Duo HPLC system.

Separation was achieved using a 50 mM sodium phosphate, 300 mM NaCl, 4% IPA, pH 6.8 mobile phase, at a flow rate of 0.2 mL/min, with an 11-minute injection time. Elution was monitored by UV absorbance at 280 nm.

Samples with concentration below 0.05 mg/mL were not run in aSEC due to lack of signal and unreliable data at lower concentrations.

Chromatograms were analyzed using Chromeleon CDS software. Peak retention times and areas were used to assess the proportion of monomeric protein as well as the presence of high molecular weight aggregates or low molecular weight degradation products.

#### C.4 Dynamic Light Scattering (DLS)

Protein particle size distribution and solution homogeneity were characterized using dynamic light scattering (DLS). This approach offers high sensitivity to early aggregate formation, allowing detection of subtle variations in conformational behavior and sample composition.

Each protein sample was loaded into standard capillaries (Cat.# PR-AC002; NanoTemper Technologies) with an approximate volume of 10 µL and analyzed using the Prometheus Panta instrument. Sample-specific details including identity, buffer conditions, and concentration were entered into the software platform (PR Panta Analysis, Version 1.9) before measurements were initiated.

DLS measurements yielded two principal values: the hydrodynamic radius (*r_H_*) and the polydispersity index (PDI). The *r_H_* reflects the average diffusion-based size of particles in solution, indicative of the protein’s physical state. The PDI is an indicator of size uniformity within the sample population; lower values suggest minimal heterogeneity and a well-behaved molecular species, while elevated values point to aggregation, misfolded structures, or the presence of particulate impurities (https://nanotempertech.com/).

#### C.5 Nano Differential Scanning Fluorimetry (nDSF)

Samples were introduced into the Prometheus Panta system (NanoTemper Technologies) using standard capillaries (Cat.# PR-AC002; NanoTemper Technologies) with a loading volume of approximately 10 µL. Prior to measurement and data acquisition, sample metadata including identification, concentration, and buffer composition were recorded using the system’s control software (PR Panta Analysis, Version 1.9).

An initial discovery scan was conducted to optimize excitation parameters by determining the appropriate light intensity tailored to the optical characteristics of each sample. Following this calibration step, thermal unfolding experiments were performed by gradually increasing the temperature from 20 °C to 95 °C at a constant ramp rate of 1 °C per minute.

Fluorescence signals were captured simultaneously at 330 nm and 350 nm throughout the scan to monitor temperature-dependent changes in protein conformation. Structural changes during heating lead to changes in the environment of aromatic residues, primarily tryptophans, causing a measurable shift in their emission profile. In rare instances where tryptophans are absent, tyrosine fluorescence contributed to the signal.

The ratio of fluorescence intensity at 350 nm to 330 nm (F350/F330) was plotted against temperature. The first deviation from baseline was defined as the onset temperature (*T*_onset_). The point of maximum slope, derived from the first derivative of the fluorescence ratio curve, was used to determine the melting temperature (*T_m_*).

#### C.6 Affinity-capture Self-interaction Nanoparticle Spectroscopy (AC-SINS)

To evaluate the self-interaction potential of protein candidates, the AC-SINS assay was carried out following established protocols [Jain et al., 2017].

Gold nanoparticles (Cat.#15705; Ted Pella Inc.) were coated with a mixture of antibodies: 80% anti-human IgG Fc-specific goat antibody (Cat.#109-005-098; Jackson ImmunoResearch) and 20% non-specific goat polyclonal antibody (Cat.#005-000-003; Jackson ImmunoResearch).

Protein samples were incubated with the functionalized nanoparticles for 2 hours at room temperature. Spectral absorbance was recorded using an Agilent Biotech Epoch Microplate Spectrophotometer, and shifts in peak wavelength relative to control samples were used to quantify self-association.

#### C.7 Hydrophobic Interaction Chromatography (HIC)

Hydrophobic interaction chromatography (HIC) was performed to evaluate the relative hydrophobicity of antibody samples, which can be indicative of developability risks such as aggregation and nonspecific interactions [Estep et al., 2015, Jain et al., 2017]. The assay was conducted using a Proteomix HTC Butyl-NP5 column, 4.6 × 100 mm (Cat. #431NP5-4610; Sepax Technologies).

Antibody samples (5 *µ*g at 1 mg*/*mL) were diluted with mobile phase A (1.8 M ammonium sulfate (Cat. #J64419.A3; Thermo Scientific) and 0.1 M sodium phosphate (Cat. #S373-500; Fisher Scientific; Cat. #S468-500; Fisher Scientific), pH 6.5) to achieve a final ammonium sulfate concentration of ∼ 1 M prior to injection. Chromatographic separation was performed using a linear gradient elution over 20 minutes, transitioning from mobile phase A to mobile phase B (0.1 M sodium phosphate, pH 6.5) at a flow rate of 1.0 mL*/*min. UV absorbance was monitored at 280 nm using an Agilent 1260 Infinity II HPLC system.

Retention times were determined using Agilent ChemStation OpenLab CDS software. Proteins with greater hydrophobicity exhibited longer retention times due to extended interaction with the hydrophobic stationary phase.

#### C.8 BVP ELISA

The assay was adapted from Jain et al. [Jain et al., 2017]. Baculovirus particles (Cat.# E3001; Medna Scientific) were diluted 1:200 in 50 mM sodium carbonate buffer (pH 9.6) and coated onto ELISA plates overnight at 4 °C.

Plates were blocked with PBS containing 2% BSA, followed by washing. Test antibodies (100 µg/mL) were added and incubated for 1 hour. After washing, HRP-conjugated secondary antibodies were added, followed by TMB substrate.

The reaction was stopped with 2 M sulfuric acid, and absorbance was measured at 450 nm. Values were normalized to control wells to compute BVP scores.

### D Experimental Data

#### D.1 Protein Targets and Design Sequences

**Table 3:** Summary of recombinant target proteins used in this study, including their corresponding UniProt identifiers, commercial source (Sino Biological), catalog numbers, and purification tags.

| Target | Assay valency | Material (catalog) | Oligomeric state | Tag | UniProt |
| --- | --- | --- | --- | --- | --- |
| METTL16 | Monovalent | Sino · M336-380G | Monomer | GST | Q86W50 |
| S100B | Bivalent | Sino · 10181-H07E | Homodimer | His | P04271 |
| ORM2 | Monovalent | Sino · 13299-H08H | Monomer | His | P19652 |
| MZB1 | Bivalent | Sino · 15682-H02H | Monomer | Fc | Q8WU39 |
| AMBP | Bivalent | Sino · 13141-H05H1 | Homodimer | Fc | P02760 |
| SORCN | Bivalent | Sino · 14547-H20B | Homodimer | His+GST | P30626 |
| SAE1 | Monovalent | Sino · 13921-HNCB | Monomer | Untagged | Q9UBE0 |
| PHYH | Monovalent | Sino · 13368-HNAE | Monomer | Untagged | O14832 |
| NTM1A | Bivalent | Sino · 11222-HNCE | Homodimer | Untagged | Q9BV86 |
| PMVK | Monovalent | Sino · 14583-H07E | Monomer | His | Q15126 |
| RFK | Monovalent | Sino · 13881-H07E | Monomer | His | Q969G6 |
| ONCM | Monovalent | Sino · 50112-MNAE | Monomer | Untagged | P53347 |
| HNMT | Monovalent | Sino · 13096-H09E | Monomer | GST | P50135 |
| 1433E | Bivalent | Sino · Y75-30BH | Homodimer | His | P62258 |
| LEP | Bivalent | Sino · 10221-HNAE | Monomer | Untagged | P41159 |
| UBC9 | Monovalent | Sino · 13205-HNCE | Monomer | Untagged | P63279 |
| SOMA | Monovalent | Sino · 16122-HNCE | Homodimer | Untagged | P01241 |
| 1433B | Bivalent | Sino · Y71-30H | Homodimer | His | P31946 |
| GM2A | Monovalent | Sino · 13246-H08B | Monomer | His | P17900 |
| IDI2 | Monovalent | Sino · 15292-H07E | Monomer | His | Q9BXS1 |

**Table 4:**
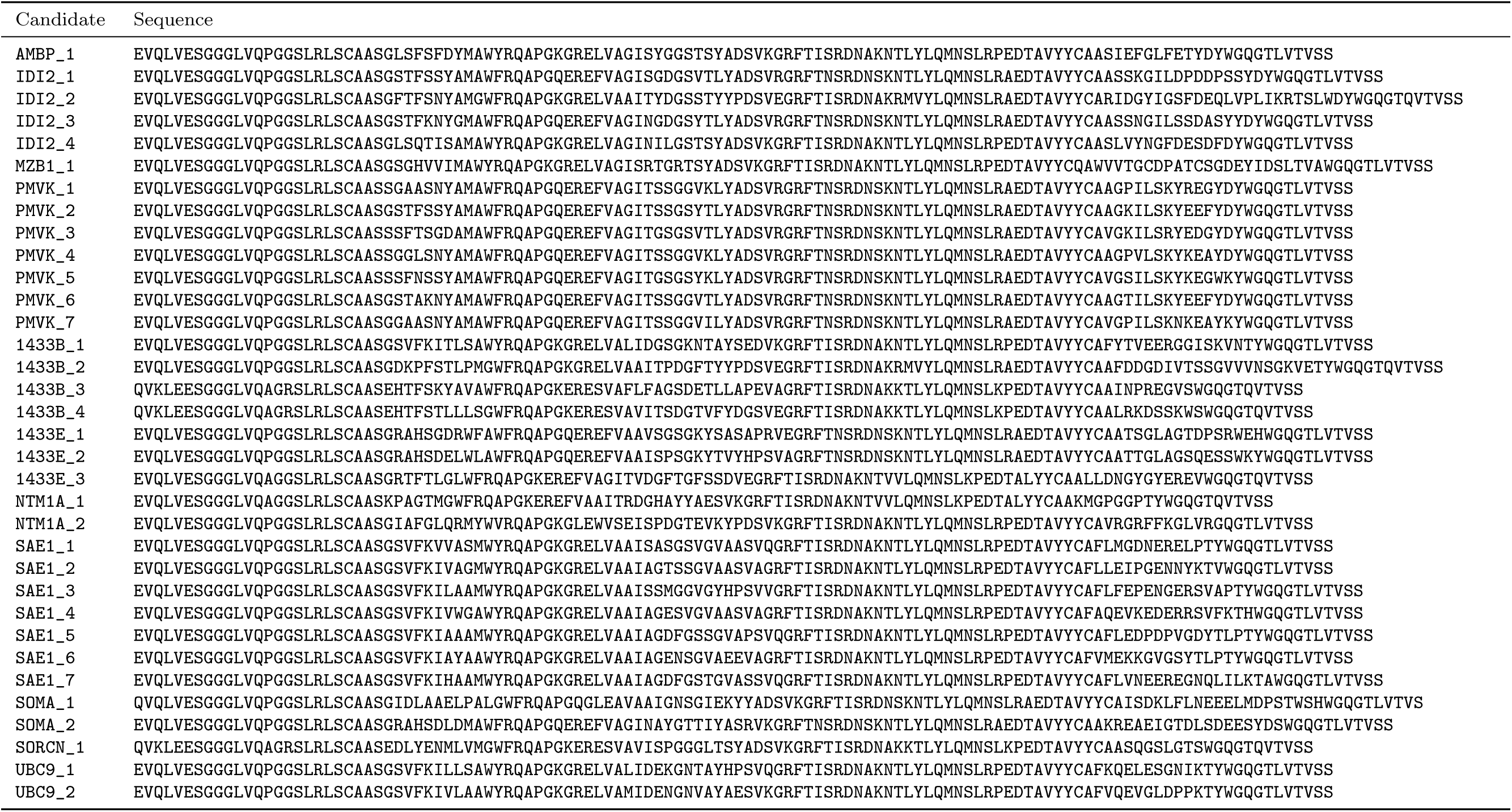
VHH sequences: 13 confirmed binders from the BoltzGen low-homology targets and 21 screening hits from Chai targets. Candidate IDs are <target>_<rank>.

#### D.2 Sensorgrams

This section provides the BLI/SPR sensorgrams for the 13 confirmed nanobody binders on the low-homology targets and 21 nanobody screening hits on Chai-2 targets. Designs are referred to by anonymized identifiers. For each binder we show the forward Adaptyv measurement(s) together with the orthogonal flipped-format assay used to confirm binding; where available we also include an independently sourced (Sino) antigen measurement.

##### D.2.1 BoltzGen low-homology targets

**Figure 8:**
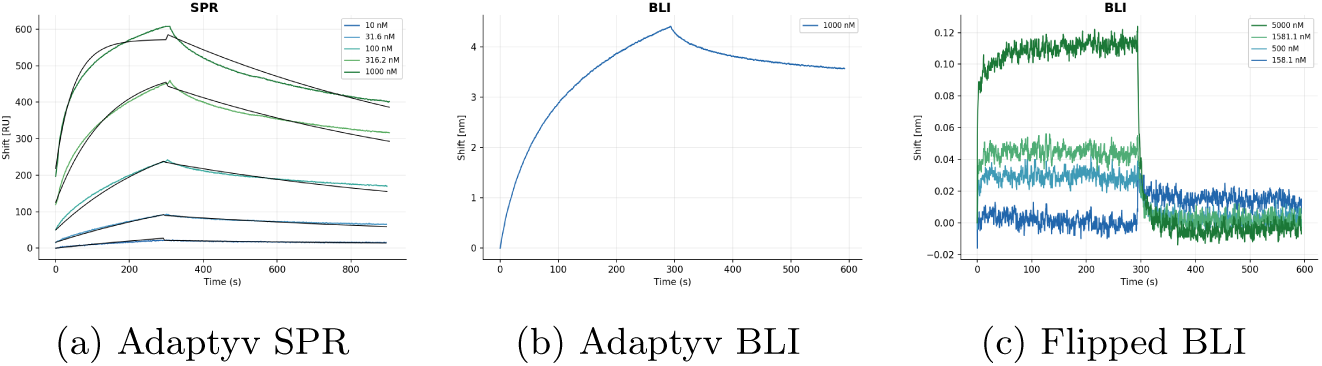
AMBP_1. Forward Adaptyv measurement(s) shown with the orthogonal flipped-format assay used to confirm binding.

**Figure 9:**
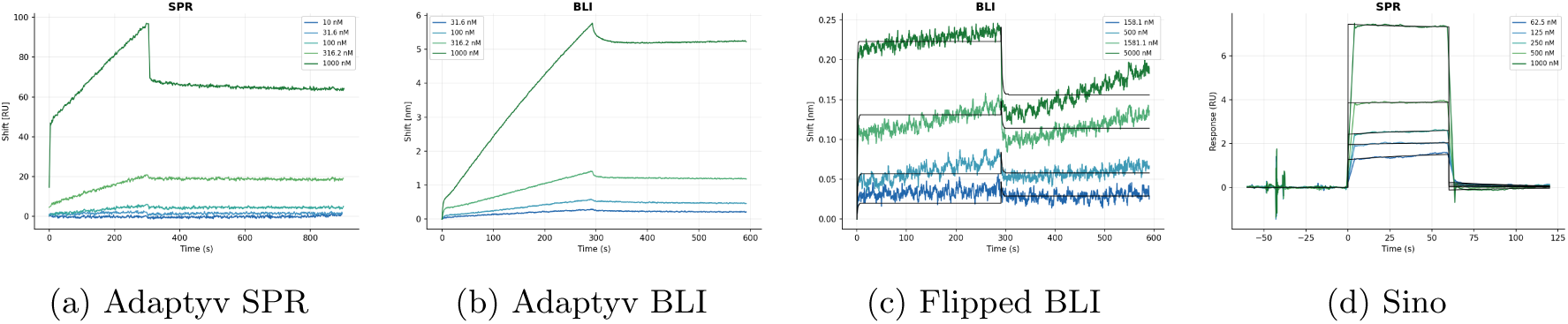
IDI2_1. Forward Adaptyv measurement(s) shown with the orthogonal flipped-format assay used to confirm binding, and an independently sourced (Sino) antigen measurement.

**Figure 10:**
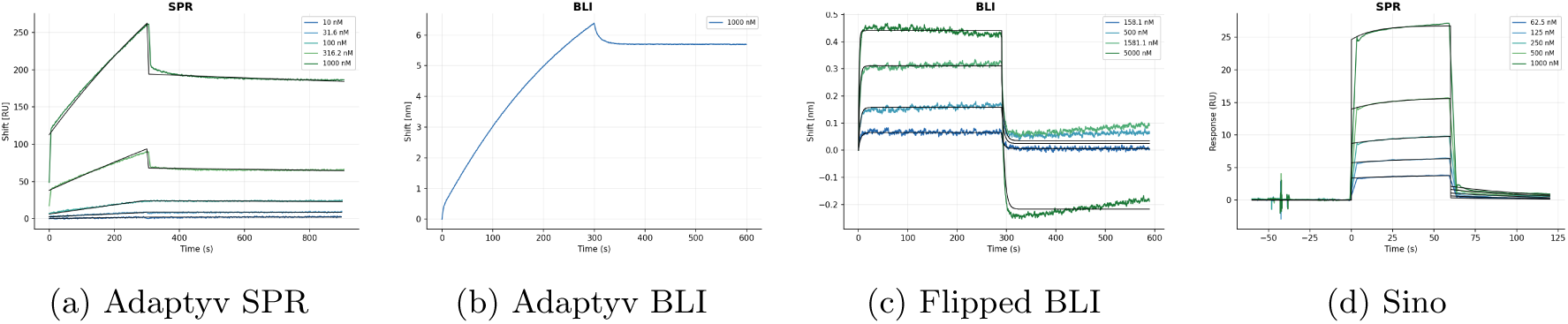
IDI2_2. Forward Adaptyv measurement(s) shown with the orthogonal flipped-format assay used to confirm binding, and an independently sourced (Sino) antigen measurement.

**Figure 11:**
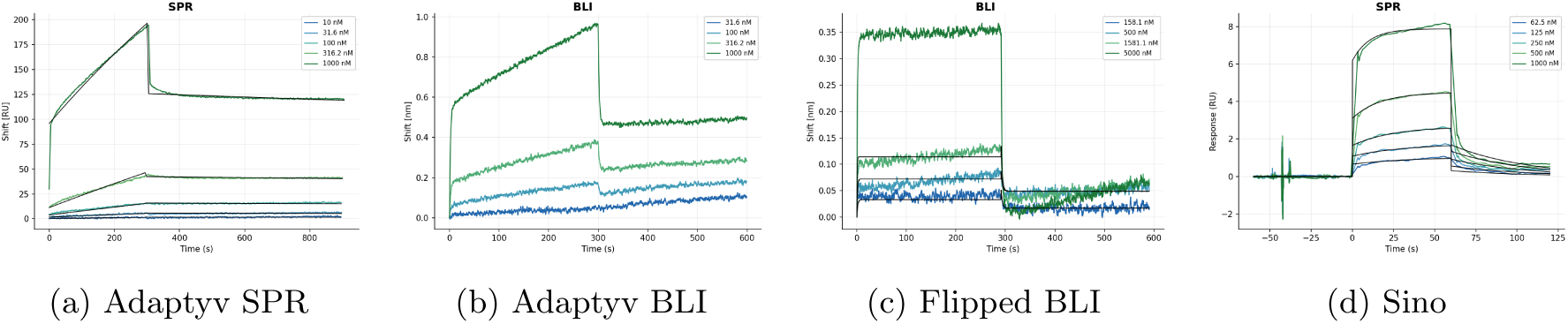
IDI2_3. Forward Adaptyv measurement(s) shown with the orthogonal flipped-format assay used to confirm binding, and an independently sourced (Sino) antigen measurement.

**Figure 12:**
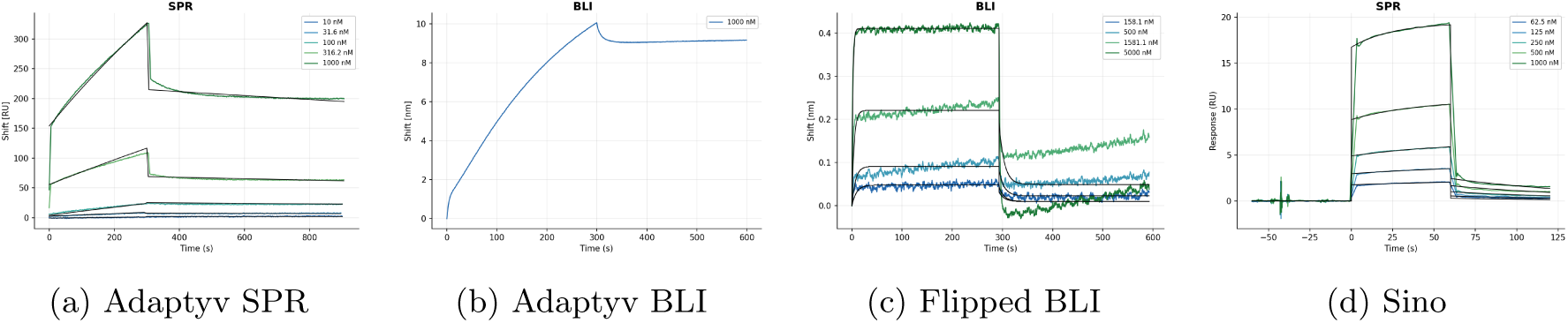
IDI2_4. Forward Adaptyv measurement(s) shown with the orthogonal flipped-format assay used to confirm binding, and an independently sourced (Sino) antigen measurement.

**Figure 13:**
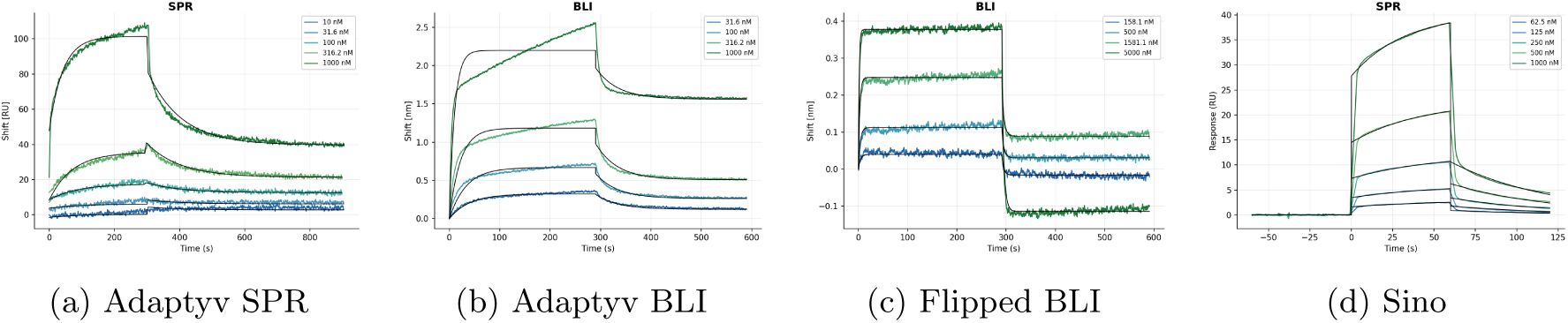
PMVK_1. Forward Adaptyv measurement(s) shown with the orthogonal flipped-format assay used to confirm binding, and an independently sourced (Sino) antigen measurement.

**Figure 14:**
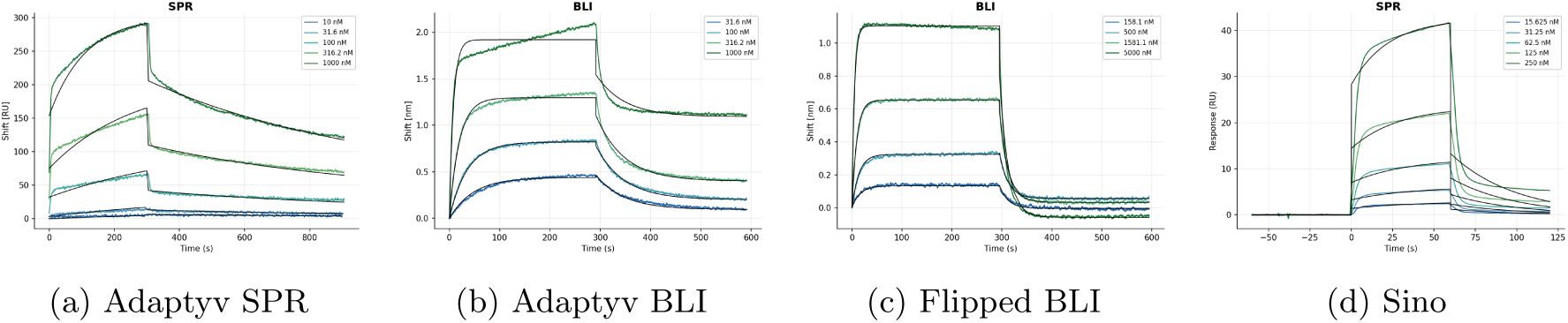
PMVK_2. Forward Adaptyv measurement(s) shown with the orthogonal flipped-format assay used to confirm binding, and an independently sourced (Sino) antigen measurement.

**Figure 15:**
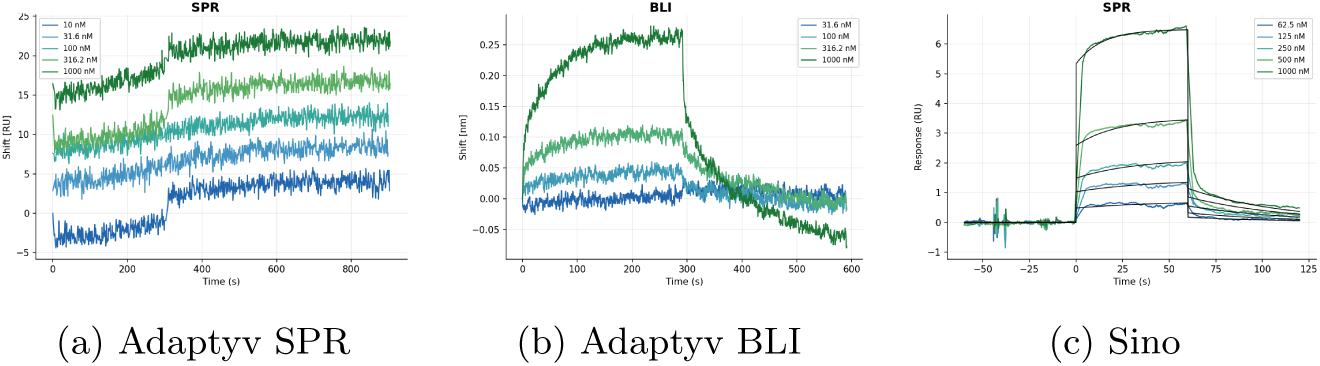
PMVK_3. Forward Adaptyv measurement(s) shown with the independently sourced (Sino) antigen measurement.

**Figure 16:**
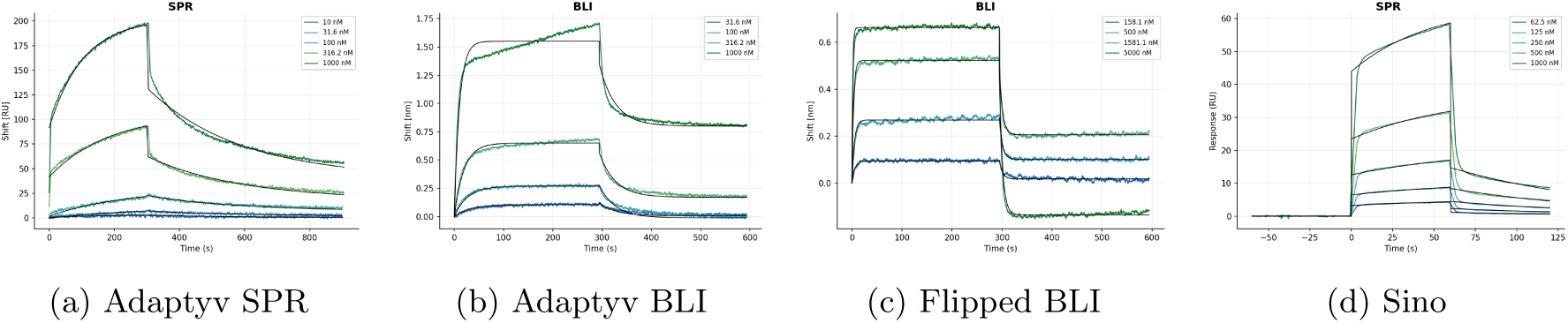
PMVK_4. Forward Adaptyv measurement(s) shown with the orthogonal flipped-format assay used to confirm binding, and an independently sourced (Sino) antigen measurement.

**Figure 17:**
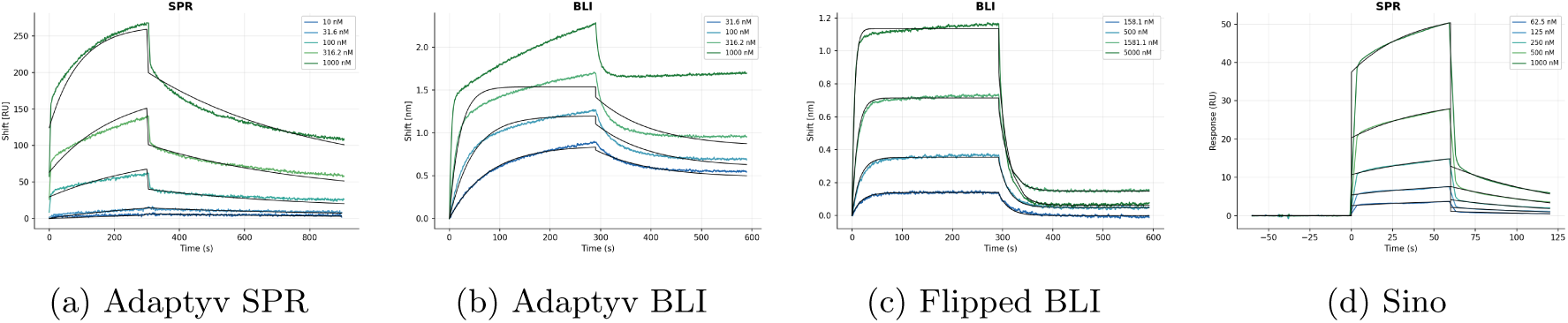
PMVK_5. Forward Adaptyv measurement(s) shown with the orthogonal flipped-format assay used to confirm binding, and an independently sourced (Sino) antigen measurement.

**Figure 18:**
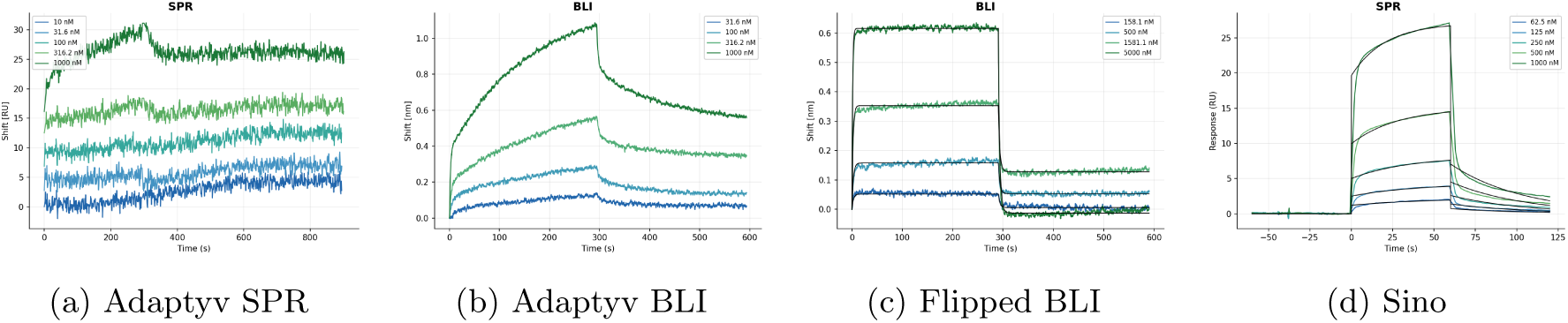
PMVK_6. Forward Adaptyv measurement(s) shown with the orthogonal flipped-format assay used to confirm binding, and an independently sourced (Sino) antigen measurement.

**Figure 19:**
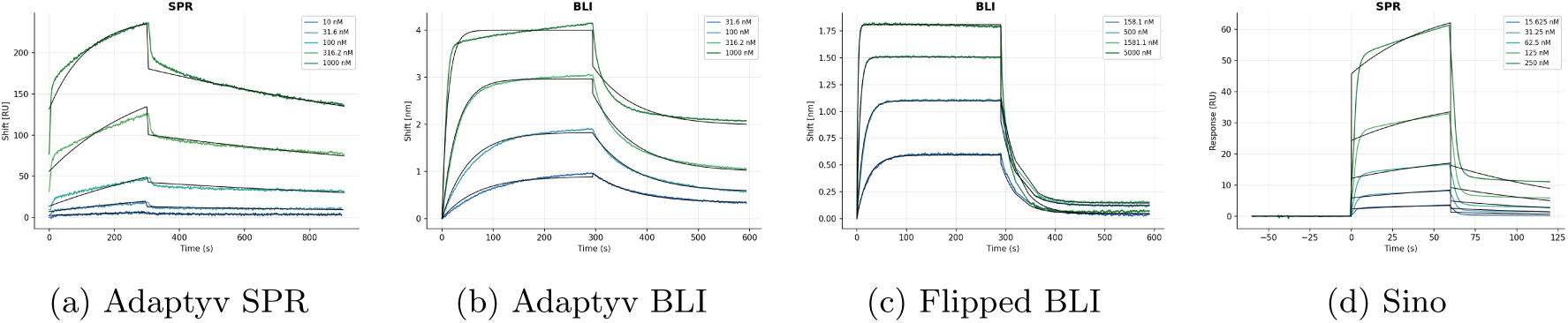
PMVK_7. Forward Adaptyv measurement(s) shown with the orthogonal flipped-format assay used to confirm binding, and an independently sourced (Sino) antigen measurement.

**Figure 20:**
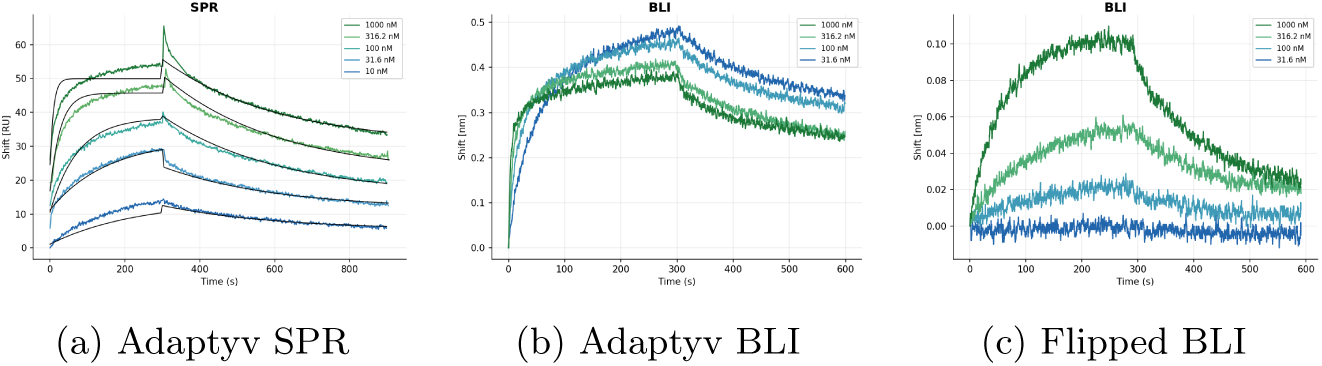
MZB1_1. Forward Adaptyv measurement(s) shown with the orthogonal flipped-format assay used to confirm binding.

##### D.2.2 Chai-2 panel

This section provides the SPR/BLI sensorgrams for the Chai-2 designs, grouped by target with one figure per target.

**Figure 21:**
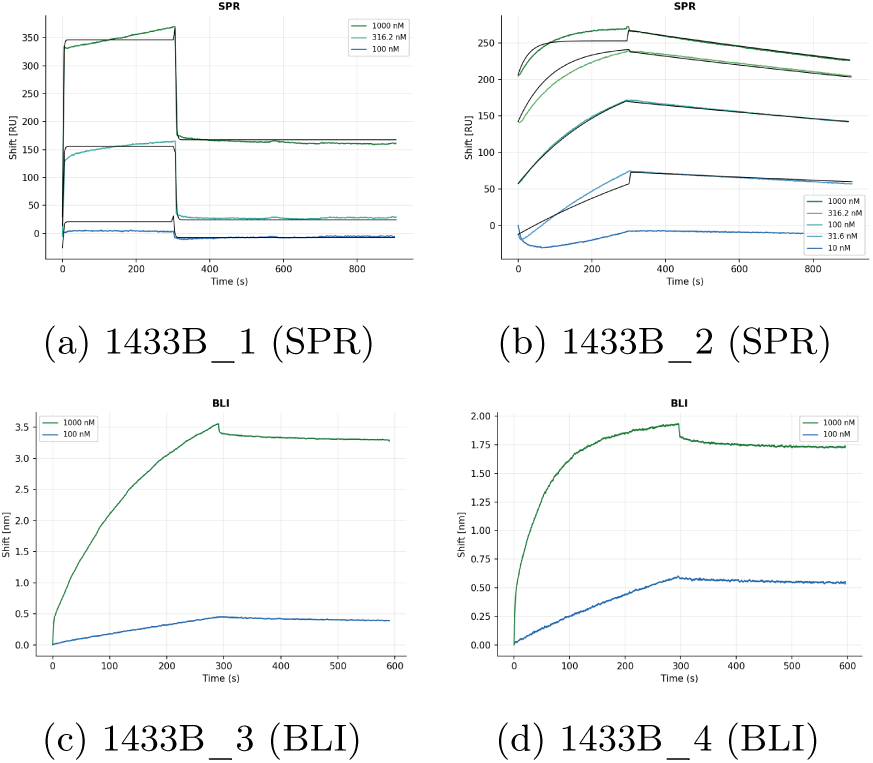
14-3-3*β* (1433B) SPR/BLI sensorgrams for the Chai-2 designs against 14-3-3*β* (1433B).

**Figure 22:**
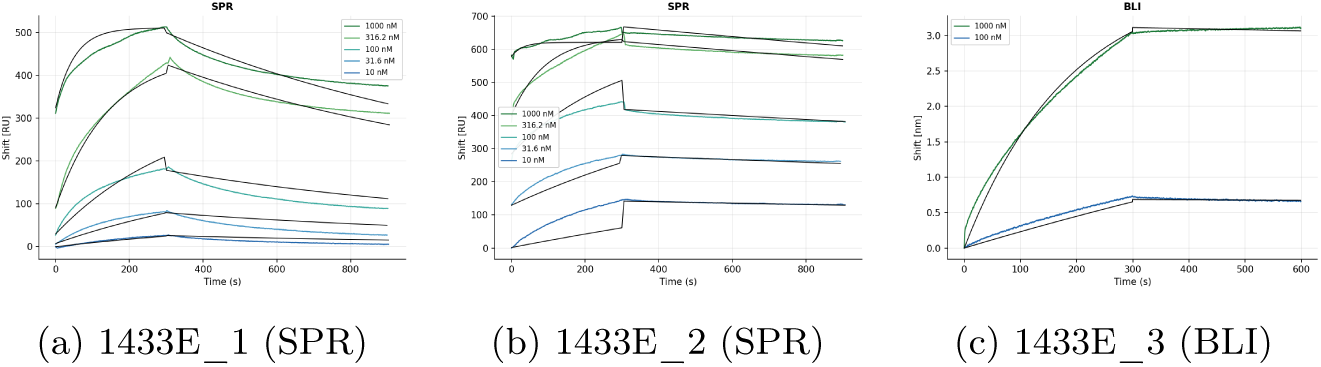
14-3-3*ε* (1433E). SPR/BLI sensorgrams for the Chai-2 designs against 14-3-3*ε* (1433E).

**Figure 23:**
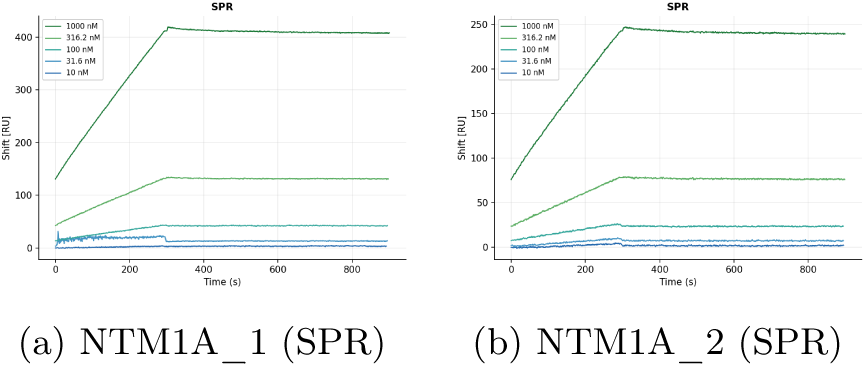
NTM1A. SPR/BLI sensorgrams for the Chai-2 designs against NTM1A.

**Figure 24:**
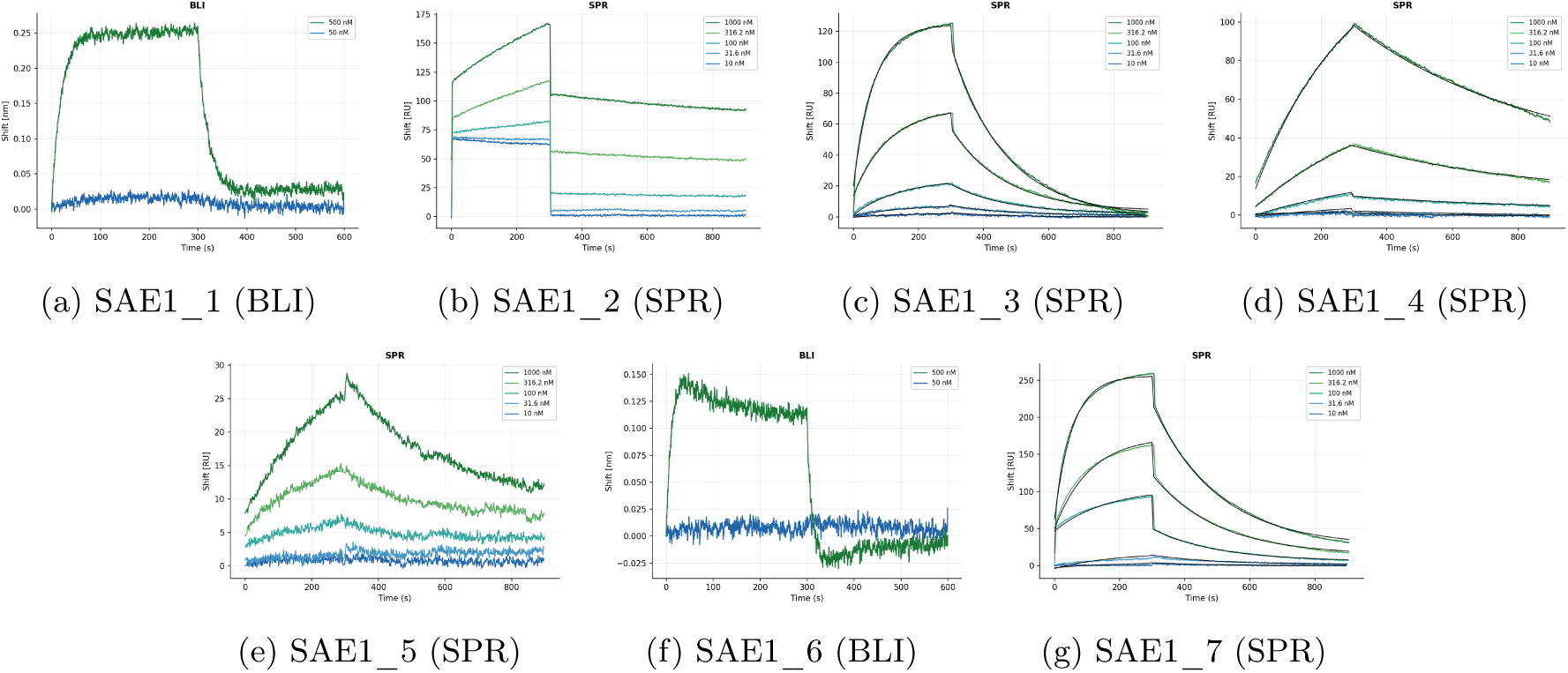
SAE1. SPR/BLI sensorgrams for the Chai-2 designs against SAE1.

**Figure 25:**
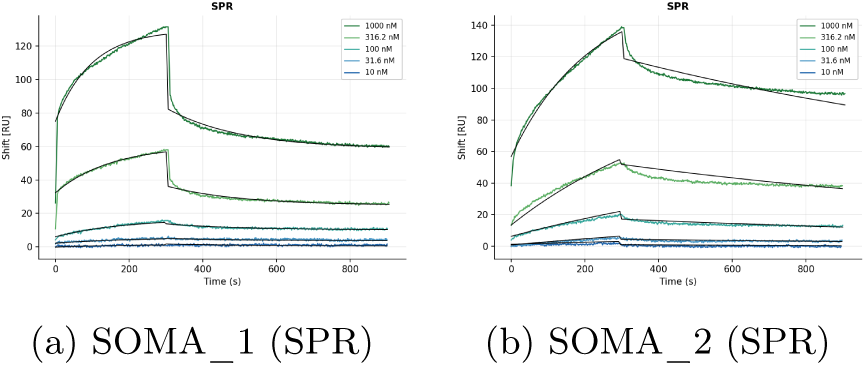
SOMA. SPR/BLI sensorgrams for the Chai-2 designs against SOMA.

**Figure 26:**
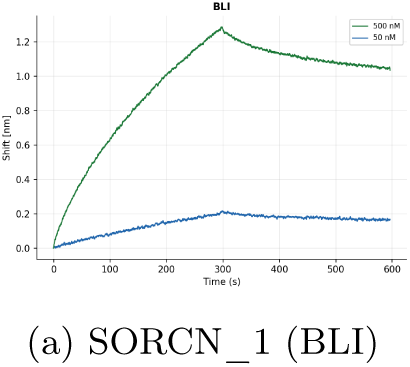
SORCN. SPR/BLI sensorgrams for the Chai-2 designs against SORCN.

**Figure 27:**
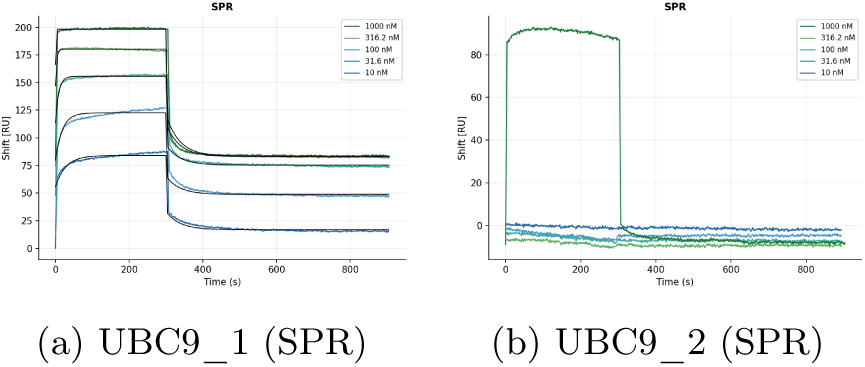
UBC9. SPR/BLI sensorgrams for the Chai-2 designs against UBC9.

### E Boltz API Usage: Protein Binder Design

Full details and instructions for implementation of the Boltz API can be found online: https://api.boltz.bio/docs/guides/protein-design

API Keys must be generated prior to launching a request: https://api.boltz.bio/docs/guides/getting-started

A *de novo* nanobody binder design run takes the target as a 3D structure template (an mmCIF file, base64-encoded) rather than a sequence. The chain_selection names the target chain and the residues exposed as the design surface (crop_residues, 0-indexed); the binder_specification requests Boltz’s curated nanobody, excluding the *N* -glycosylation sequons (*N* -*X*-*S/T*) from the binder sequence. num_proteins sets the number of designs (e.g. 60,000).

#### E.1 Example API Call

Following is an example API input for a nanobody-design run against 1433B (14-3-3*β*), with the target structure fetched directly from the RCSB PDB (entry 6A5Q) and designed against the whole target chain.

**Listing 1:**
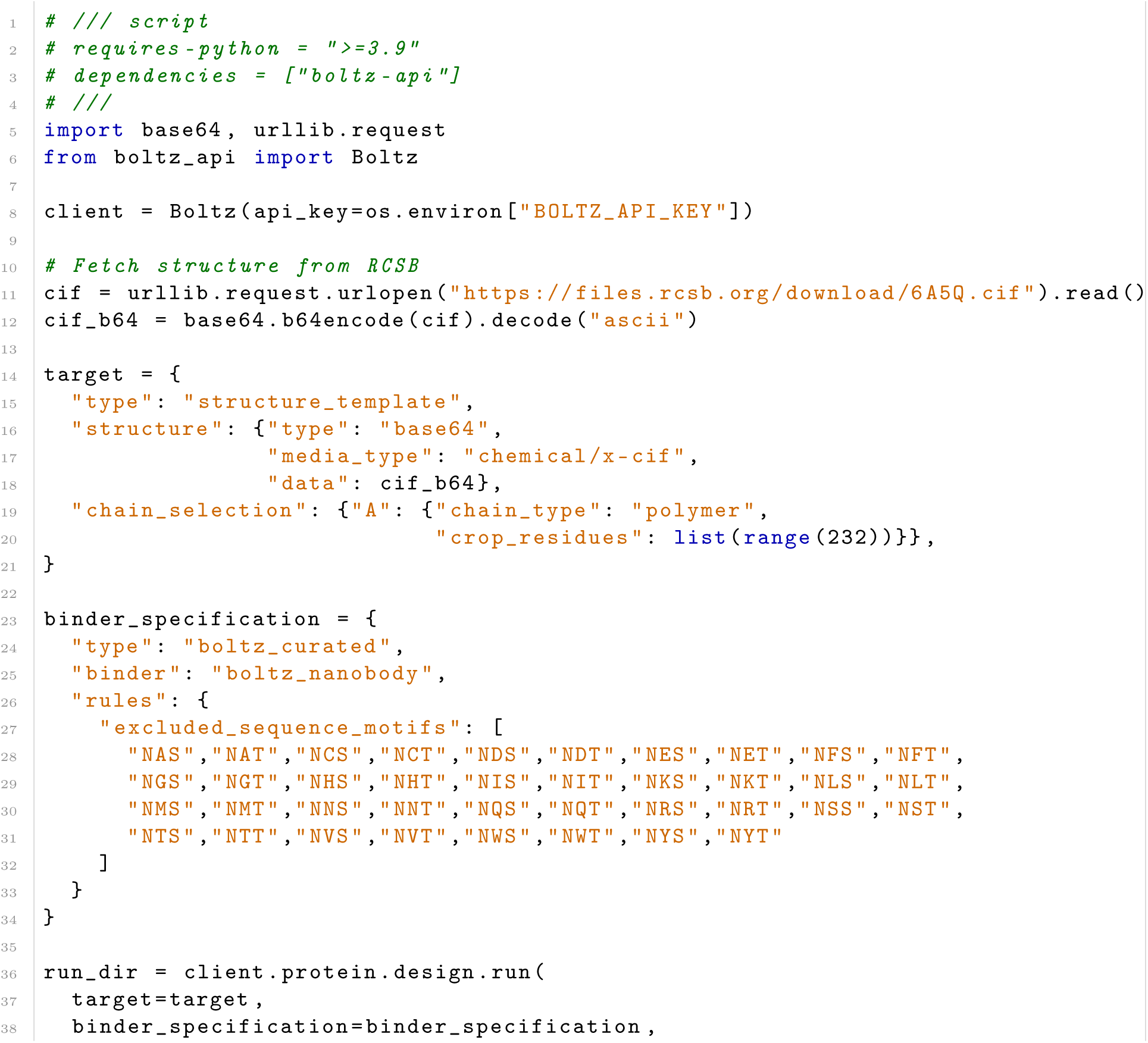
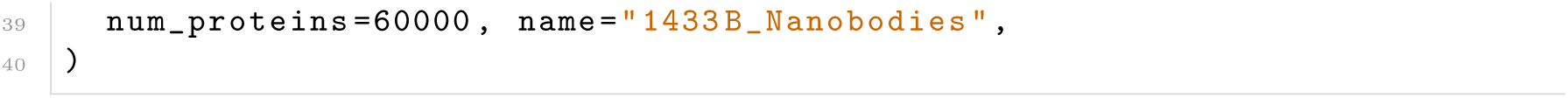
Submitting a nanobody-design run against 1433B (PDB 6A5Q) with the Boltz API.

#### E.2 Target specific inputs

The PDB and chain IDs per target are listed below. All structures are fetched from the RCSB PDB by entry ID. The chain is the Boltz API’s chain label, *not* the PDB author chain. The API renumbers chains A, B, C... in the order they appear in the file. No binding site is specified.

**Listing 2:**
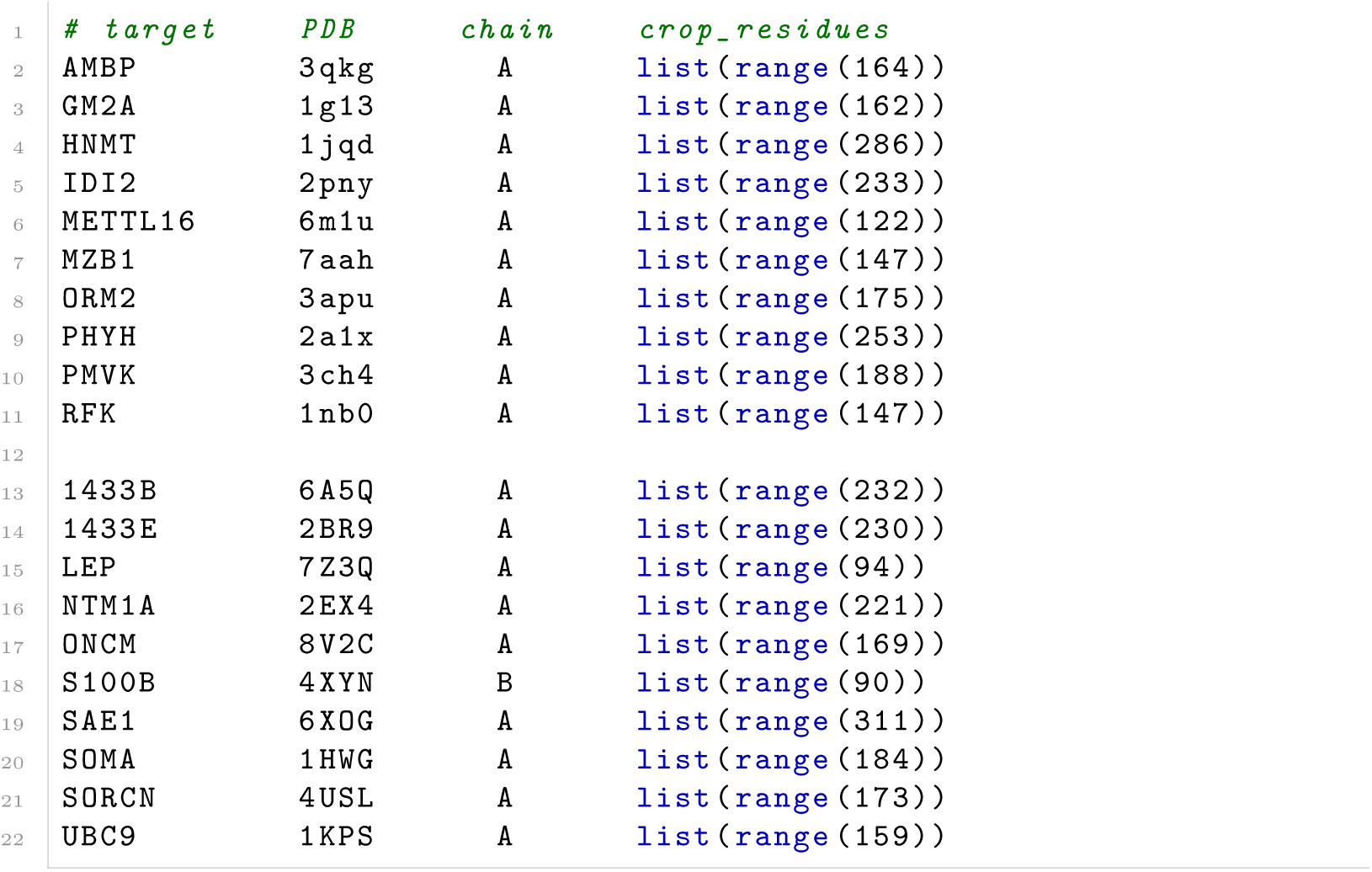
Target PDB and chain IDs.

## Footnotes

1 For instance, the target could stick to the surface itself rather than the design that is attached to the plate (which is an issue that cannot be corrected for via a reference channel if the design attachment process altered the plate itself).

